# Climatic niche conservatism in a clade of disease vectors (Diptera: Phlebotominae)

**DOI:** 10.1101/2022.01.05.475052

**Authors:** Emmanuel R.R. D’Agostino, Rafael Vivero, Luis Romero, Eduar Bejarano, Allen H. Hurlbert, Aaron A. Comeault, Daniel R. Matute

## Abstract

Sandflies of the family Psychodidae show notable diversity in both disease vector status and climatic niche. Some species (in the subfamily Phlebotominae) transmit *Leishmania* parasites, responsible for the disease leishmaniasis. Other Psychodidae species do not. Psychodid species’ ranges can be solely tropical, confined to the temperate zones, or span both. Studying the relationship between the evolution of disease vector status and that of climatic niche affords an understanding not only of the climate conditions associated with the presence and species richness of *Leishmania* vectors, but also allows the study of the extent to which psychodid flies’ climatic niches are conserved, in a context with implications for global human health. We obtained observation site data, and associated climate data, for 223 psychodid species to understand which aspects of climate most closely predict distribution. Temperature and seasonality are strong determinants of species occurrence within the clade. We built a mitochondrial DNA phylogeny of Psychodidae, and found a positive relationship between pairwise genetic distance and climate niche differentiation, which indicates strong niche conservatism. This result is also supported by strong phylogenetic signals of metrics of climate differentiation. Finally, we used ancestral trait reconstruction to infer the tropicality (i.e., proportion of latitudinal range in the tropics minus the proportion of the latitudinal range in temperate areas) of ancestral species, and counted transitions to and from tropicality states, finding that tropical and temperate species respectively produced almost entirely tropical and temperate descendant species, a result consistent for vector and non-vector species. Taken together, these results imply that while vectors of *Leishmania* can survive in a variety of climates, their climate niches are strongly predicted by phylogeny.

## INTRODUCTION

Climatic variables affect how individual species are distributed across the globe as well as patterns of diversification for clades of species (Haffer 1997; Barnagaud et al. 2012; Ali and Aitchison 2014). A notable pattern that has emerged from studying the climatic niche distribution across related species is that related species tend to have similar climatic niches. This pattern, known as niche conservatism, has been reported for multiple taxa across the tree of life including frogs (Wiens et al. 2006), mammals (Cooper et al. 2011), and angiosperms (Kerkhoff et al. 2014). Niche conservatism in climatic tolerances specifically may have implications for speciation, and how clades have expanded geographically over time. In particular, this pattern has been invoked as a potential explanation for the higher species diversity in the tropics because tropical species might not be able to evolve the ability to colonize temperate areas of the planet (Wiens and Donoghue 2004). Niche conservatism also has important implications for how species are expected to respond to global climate change because increases in global temperature will likely affect tropical and temperate clades differently (Wiens and Graham 2005).

Sandflies of the genera *Lutzomyia* and *Phlebotomus* are the only known vectors of *Leishmania*, a trypanosome parasite responsible for the disease leishmaniasis. *Lutzomyia* (Diptera: Psychodidae) is endemic to the New World and encompasses over 400 species (Young and Duran 1994). *Phlebotomus* (Diptera: Psychodidae) is endemic to the Old World and encompasses 50 species (Lewis and Lane 1976). At least 120 species from these two genera transmit leishmaniasis. Some of the *Leishmania-vector* species and others not known to transmit *Leishmania* also transmit Bartonellosis and arboviral infections (Caceres et al. 1997; Villaseca et al. 1999; Ulloa et al. 2018). For leishmaniasis alone, more than 12 million people are infected and over 2 million new cases are reported annually (Desjeux 2004; Karimkhani et al. 2016; Bailey et al. 2017). Even though species within two other genera in the family (*Sergentomyia* and *Warileya*, (Lawyer et al. 1990; Mukherjee et al. 1997; Campino et al. 2013; Kanjanopas et al. 2013; Moreno et al. 2015)) can be infected with *Leishmania*, they do not transmit the parasite. All other species in the family are known as moth-flies, some of which can cause human myasis (i.e., infection of skin tissue with larvae (Sarkar et al. 2018; Pijáček and Kudělková 2020)) and are commonly human commensals (Sparkes and Anderson 2010).

Despite the negative impacts that *Leishmania*-vectoring species (subfamily Phlebotominae) can have on human well-being, the potential drivers of geographic distribution remain highly unexplored for this group. While there are significant gaps in our knowledge of the genetics and evolutionary history of the Psychodidae, the geographic ranges of the genera in the family, in particular *Lutzomyia* and *Phlebotomus*, have been extensively characterized. In non-psychodid taxa, previous ecological niche modelling efforts have predicted the inferred range of individual vector species (Oliveira et al. 2017), and whether certain vectors are likely to expand their geographical range (Cromley 2003; Bouzid et al. 2014; Kamal et al. 2018). However, to our knowledge, the efforts to reconstruct the relationships between psychodid species have focused on taxonomic classifications and have not addressed how climate tolerance traits have evolved in the clade. For example, no study has addressed whether the distributions of these vectors are influenced by their phylogenetic relationships; namely, whether closely related species of vectors show similar geographic distributions and climatic niches or, on the contrary, have experienced climatic niche shifts over time. This is an important question because understanding the environmental variables associated with species occurrence sheds light on the potential drivers of niche evolution but also allows prediction concerning whether their ranges will expand over time (Pearman et al. 2008, 2010).

Macroecological analyses that combine environmental data with species occurrence records can reveal the extent of climatic variation across the geographic range of a species group (Diniz-Filho and Bini 2008; Keith et al. 2012). Coupled with phylogenetic analyses, macroecological data can reveal the extent of climate niche evolution in a group. In the case of vectors, these analyses can reveal whether the clinical importance of these species is likely to increase as they expand their range. Despite significant gaps in our knowledge of the genetics and evolutionary history of the Psychodidae, the geographic ranges of the genera in the family, in particular *Lutzomyia* and *Phlebotomus*, have been studied for decades. Extensive collections exist of these vectors and previous ecological niche modelling efforts have predicted the inferred range of individual vector species (Oliveira et al. 2017), and whether certain vectors are likely to expand their geographical range (Cromley 2003; Bouzid et al. 2014; Kamal et al. 2018).

In this study, we used geolocated occurrence data to determine the primary axes of climatic variation that distinguish geographic ranges of *Lutzomyia, Phlebotomus* and related genera. We find extensive variation in the climatic niche among genera within the Psychodidae and among species within the two vector genera. We find evidence that the climate niche has a strong phylogenetic signal in the family. Thermal niche differentiation between species pairs increases as divergence increases, following the expectations of niche conservatism. Temperate species are more likely to give rise to temperate species, tropical species are more likely to give rise to other tropical species, and transitions between these latitudinal zones are rare. Our work constitutes a systematic treatment of niche evolution in a family of vectors.

## METHODS

### Occurrence data

We obtained coordinates for the collections of 223 species in the Psychodidade family from Global Biodiversity Information Facility (GBIF; https://www.gbif.org/). The dataset included species from the genera *Lutzomyia* (45), *Pericoma* (24), *Phlebotomus* (26), *Psychoda* (26), *Sergentomyia* (15), *Telmatoscopus* (15), *Brumptomyia* (14), *Satchelliella* (12), *Psychodopygus* (9), *Trichomyia* (8), *Evandromyia* (6), *Clytocerus* (5), *Philosepedon* (4), *Psathyromyia* (4), *Pintomyia* (4), *Micropygomyia* (3), *Migonemyia* (2), and *Warileya* (1). The DOIs for each of the datasets are listed in Table S1. We only included the 126 extant species for which at least five georecorded locations were available.

### Estimating species’ climatic niche

Our first goal was to describe how the abiotic environment varies across sites where psychodid species have been observed. We used the collection location data described above and bioclimatic variables extracted from publicly available databases to estimate variation in the abiotic environments of Psychodidae species. For each psychodid occurrence record, we extracted four climatic variables from BIOCLIM (Booth et al. 2014) warmest-month maximum temperature, coldest-month minimum temperature, annual precipitation, seasonality of precipitation; Booth et al. 2014) plus elevation (Fick and Hijmans 2017). We chose to only consider five environmental variables to avoid overfitting, given that 70% of species had fewer than 20 occurrence records. For each of these five climate variables, we calculated 25th, 50th, and 75th percentiles of the distribution for each species. We conducted a principal component analysis (function *prcomp;* library *stats*, (R Core Team 2016)) using the 25th and 75th percentile of each trait for each species (10 total variables) in order to capture the climatic breadth of each species. However, because the eigenvectors associated with 25th versus 75th percentile were highly correlated within each climatic variable (see Results), we opted to instead use a PCA of the medians of each variable (5 total variables). To measure the extent of the differentiation along each principal component (PC) axis, we use One-Way ANOVAs where each PC was the response and the genus was the only fixed effect (function *lm*, library *‘stats’*, (R Core Team 2016)). We followed the ANOVA with Tukey’s honest difference post-hoc comparisons (function *glht*, library *‘multcomp’*, (Hothorn et al. 2020)).

### Mean latitude and tropicality index

We calculated the mean latitude of occurrence for each species as one way of characterizing geographical distribution. However, mean latitude is unable to distinguish between a range-restricted tropical species and a cosmopolitan species with an identical range centroid. As such, we additionally calculated Kerkhoff et al.’s (Kerkhoff et al. 2014) tropicality index (TI) for each species as the proportion of its latitudinal range that falls within the tropics minus the proportion of the latitudinal range that falls within the temperate zone. The index ranges between −1 (strictly temperate species) to 1 (strictly tropical species). Values of 0 correspond to species whose distribution is half-temperate and half-tropical. Following Kerkoff et al. (2014), we further assigned species to one of four distributional categories based on the tropicality index: “tropical” species with TI > 0.5 (75% or more of the range within the tropics), “semitropical” species with 0 < TI ≤ 0.5, “semitemperate” species with −0.5 < TI <= 0, and “temperate” species with TI ≤ −0.5 (75% or more of the range within the temperate zone).

### mtDNA genealogy

In spite of the rich geographical range dataset for Psychodidae, few efforts have addressed the phylogenetic relationships between species of the family. We obtained sequences of the mitochondrial locus Cytochrome Oxidase I data (*COI*) for 125 species from GenBank. All the accession numbers are listed in Table S2. Of these 125 species, 74 also had geographic information (described above, Table S1). We aligned the sequences using Clustal Omega (Sievers et al. 2011; Sievers and Higgins 2018) with the following specifications: -dealign -t --seqtype={DNA} --outfmt=phylip -v. We included the *COI* sequence for *Aedes albopictus* to serve as an outgroup and root the tree (Table S1).

We generated a phylogenetic tree for the 74 species for which we have both mtDNA and geographic information using IQTREE, and these species formed the core of our analyses for this study. We used the -m TEST option (jModelTest, (Posada 2008; Darriba et al. 2012)) for model selection. To estimate support of the branches we used SH-aLRT support (%) and ultrafast bootstrap support (%) using 1,000 replicates. We only kept nodes with 60% bootstrap support. Here, we do not aim to infer all the genealogical relationships between species in the Psychodidae, as more data than just the *COI* locus would be required for this task. The DRYAD data package (DOI:TBD) contains the log files for the run.

### Phylogenetic signal

We estimated the phylogenetic signal of the climatic niche (characterized by the PC1 score from the ordination above), the mean latitude of each species, and the tropicality index across the Psychodidae tree using two different metrics: Blomberg’s *K* (Blomberg et al. 2003), and Pagel’s *Λ*. Both metrics were calculated using the function *‘phylosig’* (library *phytools*, (Revell 2012)) with 1,000 simulations to determine if the calculated value differed from zero. Blomberg’s *K* (Blomberg et al. 2003) indicates whether the association between the tree and the trait follows the expectations under a Brownian model of evolution (i.e., the trait value changes randomly, in both direction and magnitude, over the course of evolution). If *K* equals 1, the evolution of species’ traits (climatic niche, mean latitude, tropicality index) conforms to a model of Brownian motion evolution in which trait values of descendant species diverge slowly from the ancestral value. A *K* lower than one suggests that relatives resemble each other less than expected under Brownian motion evolution and is evidence against phylogenetic niche conservatism (Cooper et al. 2010). A *K* higher than 1 suggests that close relatives are more similar than expected under Brownian motion evolution. This can be caused by phylogenetic constraints or niche conservatism (Cooper et al. 2010).

The second metric, Pagel’s *Λ*, is a measure of phylogenetic signal which estimates the extent to which the phylogenetic history of a clade is predictive of the trait distribution at the tree tips. Values of *Λ* lower than 1.0 represent traits being less similar amongst species than expected from their phylogenetic relationships. A λ equal to 1.0 suggests that traits covary with phylogeny (Pagel 1997, 1999) and is consistent with, but not diagnostic of, niche conservatism (Cooper et al. 2010).

### Climatic niche divergence

To additionally assess whether genetic divergence and climate niche differentiation were associated, we used regression analyses. Our goal was to determine whether the relationship between genetic divergence and climatic niche differentiation had a positive slope (i.e., species became more differentiated in climate niche as genetic distance increased). To score the divergence between two species in their climatic niche, we calculated the difference between their mean PC1 values (ΔPC1). We then fit a linear regression and calculated the slope of the relationship between ΔPC1 and genetic distance (function *lm*, library *stats*; (R Core Team 2016)). We bootstrapped the regression coefficients using the function *Boot* (library *simpleboot*, (Peng 2008)). These analyses are conceptually related to the calculation of phylogenetic signal because cases where the difference in the climatic niche of a pair of species increases as divergence accrues should also have a strong phylogenetic signal.

Closely related species do not behave independently and their similarity is likely to be the result of shared history (Huey et al. 2019). We therefore used two phylogenetic non-independence corrections. First, we use a variant of phylogenetic regressions in which the genetic relationships of the species are considered a random effect in a linear mixed model. We used the function *force.ultrametric* (library *‘phytools’*, (Revell 2012)) to make an ultrametric tree and the function *cophenetic* (library *‘stats’*, (R Core Team 2016)) to generate a genetic distance matrix. We used a linear mixed model in which the difference in PC1 was the response, genetic distance was a continuous variable, and the two species in the comparison were considered random effects using the function *lmer* (library ‘*lme4*’, (De Boeck et al. 2011; Bates et al. 2013)). We used the function *bootMer* (library ‘*lme4*’, (Bates et al. 2013)) to bootstrap the regression (1,000 replicates). To compare the slope of the phylogenetically-corrected and the non-corrected regressions, we used a Wilcoxon rank sum test with continuity correction (function *wilcox.test*, library *stats*; (R Core Team 2016)).

Second, we fitted a generalized linear mixed model using Markov chain Monte Carlo. We used the function *ginvn* (library *MASS*, (Venables 2002; Venables et al. 2003)) to find the generalized inverse of the (1-genetic distance) matrix as proposed by Castillo (Castillo 2017). We fitted a linear model using the package *MCMCglmm* (Hadfield 2010) in which the difference between climatic PC1 was the response, genetic distance was a predictor variable and the phylogenetic covariance matrix was a random effect. We ran five independent MCMC chains. To determine if the model converged in each of the chains, we used the function *gelman.diag* (library *coda*, (Plummer et al. 2006)). A chain was considered converged if all scale reduction factors for all variables (both fixed and random effects) were ≤1.1 for each of the two chains. We calculated the 95% confidence interval for the intercept and the slope using the function *HPDinterval* (library *coda*, (Plummer et al. 2006)).

Next, to understand the dynamics of diversification of climatic niche in Psychodid species, we fitted seven different models of trait evolution using the function *fitContinuous* (library *‘geiger’*, (Harmon et al. 2008; Pennell et al. 2014). Models varied in the tempo and mode of trait evolution and ranged from no phylogenetic signal (i.e., white noise) to different modes of evolution. The details of the seven models are described in Table 1. First, and following (Cooper et al. 2010; Wiens et al. 2010), we compared three models of trait evolution: white noise, Ornstein–Uhlenbeck (OU), and Brownian. Support for the latter two suggests evidence of niche conservatism (Cooper et al. 2010). Next, we compared all seven models using their Akaike Information Criterion (AIC) values and calculating their Akaike weights (*wAIC*) using the equation:

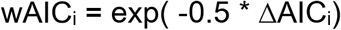

Where ΔAIC_i_ is the difference between the AIC of model *i* and the model with the lowest AIC. We used the function *aic.w* (library *phytools*, (Revell 2012)) for these calculations.

**TABLE 1.** Models of trait evolution fitted to climatic niche data in the Psychodidae family.

| **Model** | **Assumptions** | ***fitContinuous* option** |
| --- | --- | --- |
| **White-noise** | The trait values come from a single normal distribution with no covariance structure among species. | model="white" |
| **Lambda** | A model in which phylogeny predicts covariance among trait values (Pagel 1999). The model transforms the tree using a scalar, $\lambda$, that ranges between 0 (a star-like phylogeny) and 1 (the BM model, see below). Equivalent to calculating a form of phylogenetic signal ($\lambda$, see Methods). | model="lambda" |
| **Early Burst** | The rate of evolution increases or decreases exponentially through time, under the model $r_t = r_0 \times e^{(a \times t)}$, where $r_0$ is the initial rate, $a$ is the rate change parameter, and $t$ is time. | model="EB" |
| **Ornstein Uhlenbeck (OU)** | Trait evolution is best-explained by a random walk with a central tendency and an attraction strength determined by the parameter alpha, which ranges between ~0 and 2.72 (Butler and King 2004). | model="OU" |
| **Brownian motion model** | The correlation structure among trait values is proportional to the extent of shared ancestry for pairs of species (Felsenstein 1973). | model="BM" |
| **Kappa** | Character divergence is associated with speciation events in the tree (Pagel | model="kappa" |
|  | 1999). kappa ranges between ~0 and 1. |  |
| **Delta** | A model that fits relative contributions of early versus late evolution in the tree to the covariance of species trait values (Pagel 1999). Delta values larger than 1 suggest recent evolution has been relatively fast; delta lower than 1, suggest recent evolution has been comparatively slow. | model="delta" |

### Ancestral trait reconstruction and rates of transition

We inferred the climatic niche of each node in the Psychodidae tree using the *COI* gene genealogy for three proxies of climate niche: *i)* tropicality index (TI), *ii)* mean latitude, and *iii)* PC1 (described in ‘Estimating species’ climatic niche’).

We used the function *‘collapse.singles’* (library *ape*, (Paradis and Schliep 2019)) to resolve polytomies. Trees were then checked with the function *‘is.binary.tree’* (library *ape*, (Paradis and Schliep 2019)). We used the function ‘*anc.ML*’ (library *phylo*, (Revell 2012)) for the ancestral trait reconstruction of each of the three traits mentioned above with a maximum of 5,000 iterations using a Brownian movement model. (Similar runs using other models gave identical results.) Trees were drawn using the function *‘contMap’* (library *phylo*, (Revell 2012)). Finally, we used ancestral niche reconstruction to examine the rate at which species in the Psychodidae transitioned among and between the four latitudinal range categories (tropical, semitropical, semitemperate, and temperate) based on TI as described above (Kerkhoff et al. 2014). First, we inferred the latitudinal category for each node in the tree using *anc.ML* (Revell 2012). Ancestral trait reconstructions using other approaches (*anc.Bayes* and *fastAnc* (Revell 2012)) gave similar results. Then, for each latitudinal category, we selected all of the ancestral nodes inferred to be in that category, and calculated the proportion of immediately descendant nodes or tips that fell into each of the four latitudinal categories. Doing this for each category yielded 16 transition rates describing the frequency of transitions between any pair of latitudinal states.

## RESULTS

### Geographic distribution of Psychodidae genera

First, we calculated the mean latitude of 223 species in the Psychodidae family as a proxy of their distribution. We find that there are two peaks of diversity in the family, one in the tropics around the Equator and a smaller one slightly below the Arctic circle (Figure S1). Figure S2 shows the mean latitude for species within 12 genera in the Psychodidae family. The moth-fly genera *Clytocerus, Philosepedon, Satcheliella*, and *Telmatoscopus* are all of temperate distribution. The genera *Pericoma* and *Psychoda* are largely temperate but some species have a tropical distribution. Among *Leishmania*-harboring genera, some patterns are also salient. The genera *Lutzomyia* and *Brumptomyia* are mostly restricted to the tropics. *Phlebotomus* on the other hand shows high diversity in the northern subtropical region. These results indicate that different genera in the family show strong differences in their distribution and suggest the possibility of climate niche evolution among and within genera.

Given the strong differences in mean latitude between genera, we first tested for a relationship between environmental variables and occurrence records of each species within the different genera of psychodids included in our dataset. We found that the eigenvectors for the 25th and 75th percentiles’ contributions to the largest principal components were highly correlated (Pearson’s product-moment correlation, PC1: *r* = 0.993, *p* < 0.001; PC2: *r* = 0.980, *p* = 0.003; PC3: *r* = 0.999, *p* < 0.001). In light of this correlation, we opted to use the 50th percentile of each climate variable for each species distribution for all further analyses. A PCA revealed the relative importance of elevation, temperature, and temperature seasonality for occurrence. Table S3 shows the loadings for the PCA. The first three PCs explain the vast majority of the variance (93.96%) so we restricted our analyses to these PCs. All environmental variables had relatively high loadings on PC1 (56.72% of variation). Positive values on PC1 indicate locations that are relatively seasonal and cool, while more negative values are indicative of locations that are less seasonal and warmer (Figure 1). Genera differed along PC1 (LMM: *X^2^_1_=469.69*, P < 1 × 10^-10^, Figure 1B), but not PC2 (LMM: *X^2^_1_*= 8.48, P = 0.58), which is largely dominated by elevation (Table 2) and explained 18.37% of the variance. Finally, PC3 (17.36%), which is mostly influenced by amount and seasonality of precipitation, also differed among genera (*X^2^_1_*= 72.98, P < 1 × 10^-10^). Pairwise comparisons for these PCs all suggested strong differences across genera (Tables S4-S6).

**FIGURE 1.**
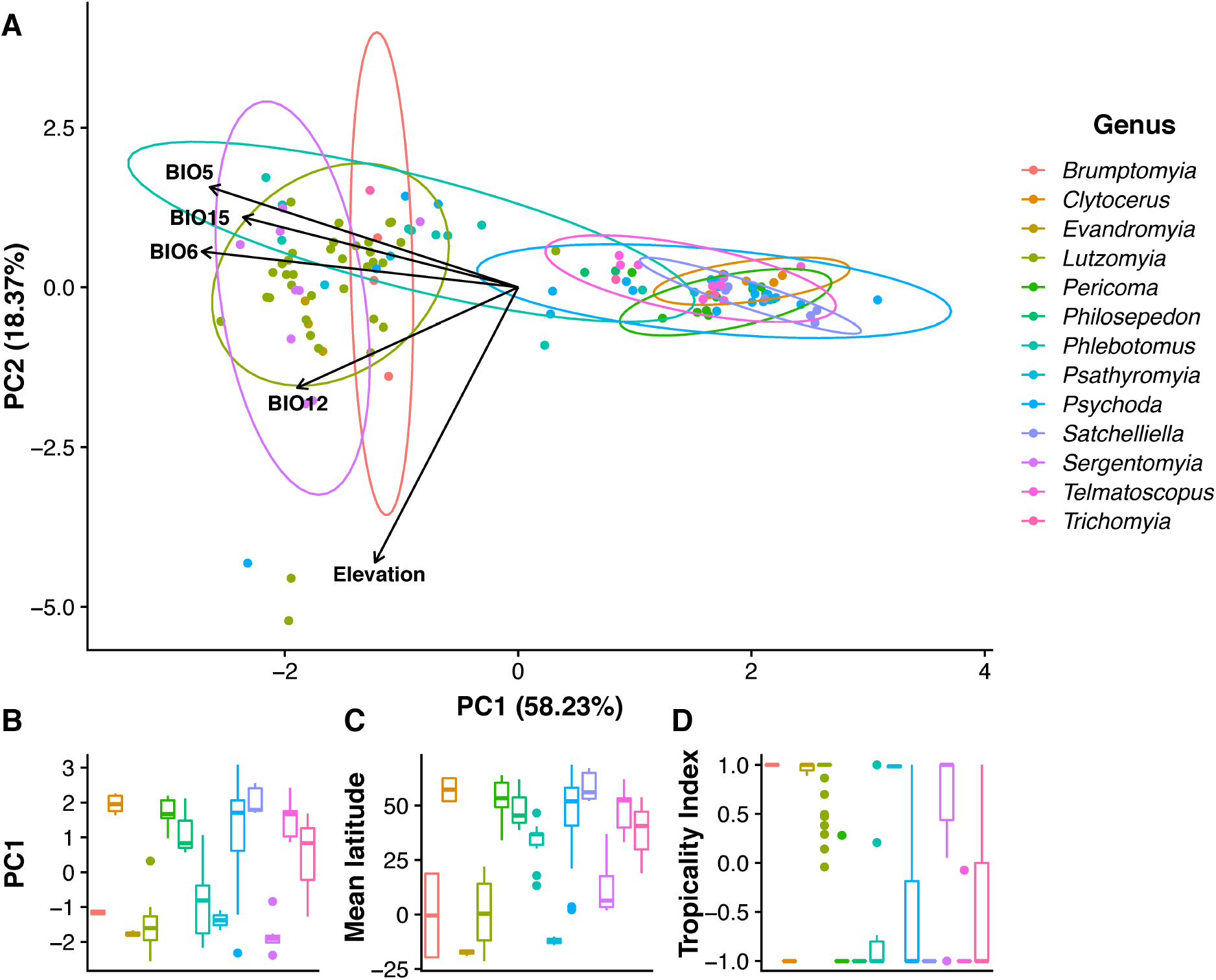
Different genera within the Psychodidae family differ from each other in their climatic niche. **A.** PCA based on the Worldclim variables suggests that the occurrence of different genera within the family are associated with climatic variables. **B.** Boxplot showing the mean species values for PC1. This PC is mostly explained by temperature and precipitation. **C.** Boxplot showing the mean species values for latitudinal distribution. **D.** Boxplot showing the mean species values for TI.

**TABLE 2.** Different proxies of climatic niche have a strong phylogenetic signal. Blomberg’s K and λ estimates for three different proxies of geographic range in the Psychodidae family.

|  | **Blomberg's K** | **$\lambda$** |
| --- | --- | --- |
| **Mean latitude** | 2.557, P=0.001 | 0.999 |
| **PC1** | 2.678, P=0.001 | 0.999 |
| **TI** | 2.175, P=0.001 | 0.999 |

### Comparative phylogenetic analyses

#### COI gene genealogy

We generated a gene genealogy based on mtDNA to get an approximation of the phylogenetic relationships within the family Psychodidae. Over 90% of branches had bootstrap support of >60% (Figure S3). Our sample contains genera from two different taxonomic subfamilies: Phlebotominae and Psychodinae. We recovered these two subfamilies as monophyletic groups (Figure 4A, Figure S3) but not all genera appear monophyletic (e.g., *Phlebotomus*). The hematophagous clade was monophyletic (*Lutzomyia, Phlebotomus, Brumptomyia* and *Sergentomyia*) but the *Leishmania* vectors were not.

#### Phylogenetic signal

We used the COI genealogy to study macroevolutionary trends of the evolutionary history of climate niche in the family. We used two complementary indices that summarise patterns of trait evolution on a phylogeny for each of our three proxies of climatic niche: TI, mean latitude, and PC1. First, we found that Blomberg’s *K* was significantly higher than 1 for each of the three metrics of geographic range (Figure 2, Table 2), indicating that the climatic niches of close relatives are more similar to each other than expected under a pure model of Brownian motion evolution (1,000 randomizations, P < 0.001; Figure 2). Second, we found that Pagel’s λ was significantly higher than 0 and lower than 1 for the three proxies (Figure 2). Broadly, these two metrics suggest that climatic descriptors of niche have a strong phylogenetic signal in the Psychodidae tree.

**FIGURE 2.**
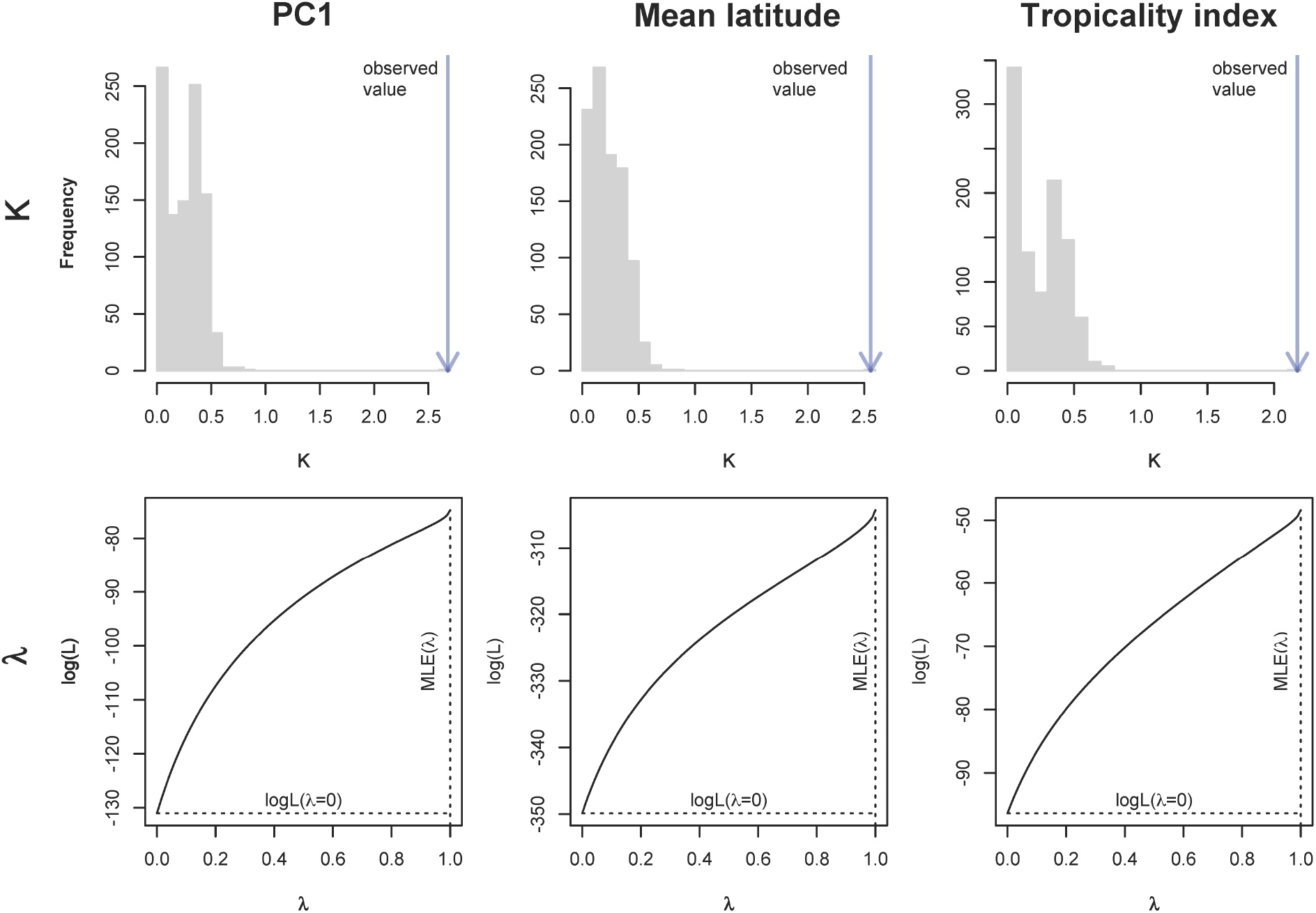
Climatic niche shows strong phylogenetic signal in the Psychodidae family. Top panels show the distribution of simulated and observed Blomberg’s K. Bottom panels show values of the maximum likelihood estimate (MLE) of λ and the maximum likelihood of the model when λ is zero. Left-side panels show metrics for PC1. Center panels show metrics for median latitude. Right-side panels show metrics for tropicality index.

Consistent with niche conservatism, we found that the genetic divergence between species pairs is positively associated with the extent of their climate niche differentiation (Figure 3; One-way ANOVA: F_1,2484_=1,754.6, P < 1 × 10^-10^). This result suggests that closely related species are the most similar in their climatic niche, and that the climate niche of Psychodidae species becomes more dissimilar as divergence increases. This result is qualitatively identical for a phylogenetically corrected dataset where the species identity are considered random effects (LMM: F_1_=1,824.06, P < 1 × 10^-10^; Figure 3, blue lines). The magnitude of the regression slope is significantly lower in the phylogenetically-corrected regression than for the non-corrected dataset (Slope_Corrected_=0.365, Slope_Non-corrected_=1.457; Wilcoxon rank sum test with continuity correction: W = 1 × 10^6^, P < 1 × 10^-10^). Finally, a MCMC-based phylogenetic correction revealed the same pattern, as the slope of the regression is also positive (95% CI = 0.0676-0.273). The results from all these analyses (i.e., a positive correlation between divergence in climatic traits and age of divergence) are consistent with our phylogenetic signal analyses, which suggest that climatic niche evolution in the species from the Psychodidae family follows the expectation of niche conservatism.

**FIGURE 3.**
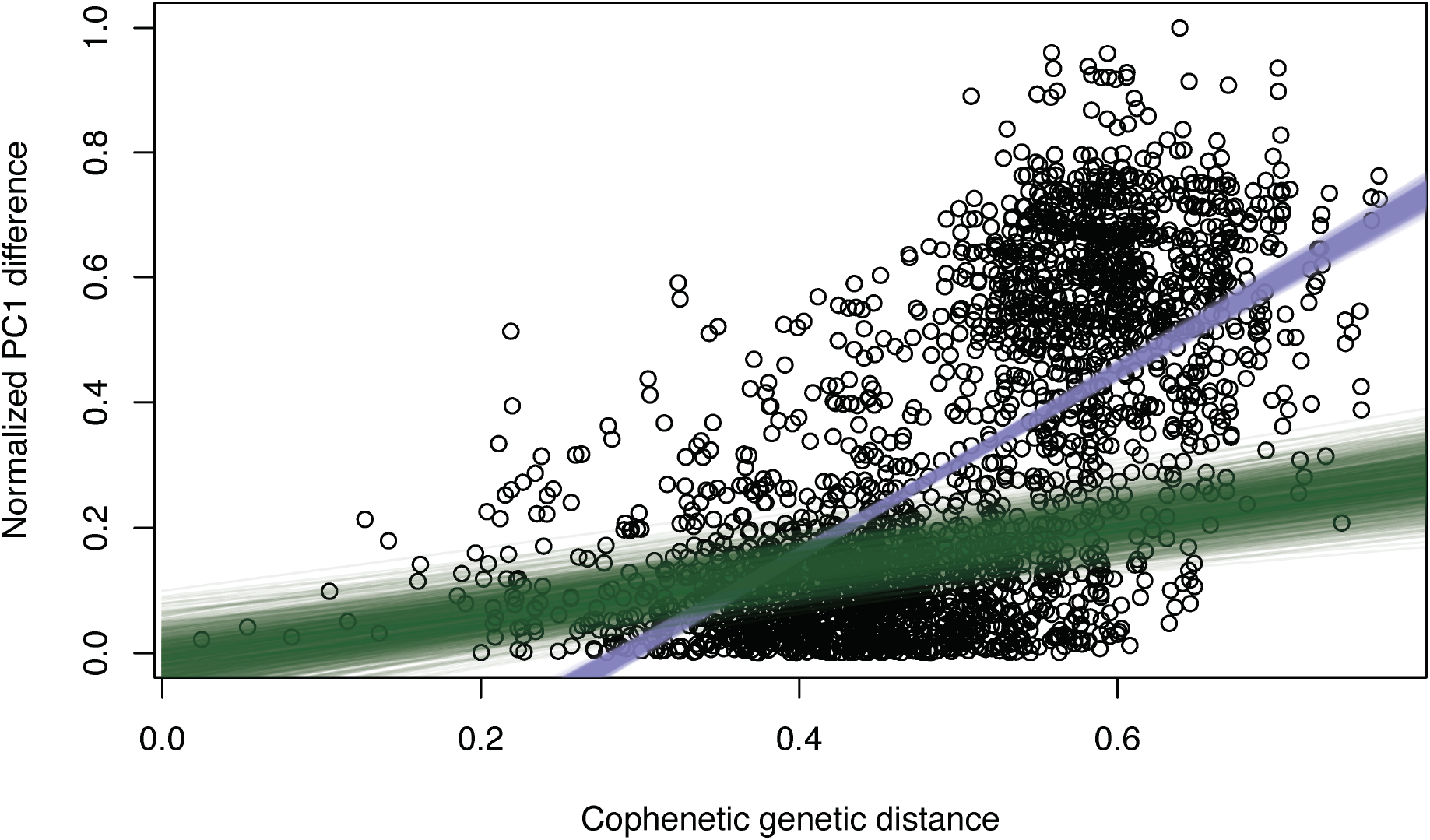
Habitat differentiation increases with genetic distance in the Psychodidae family. Blue lines show 1,000 bootstrapped linear regressions with this non-phylogenetically-corrected dataset. Green lines show 1,000 bootstrapped linear regressions with a species-identity random effect to account for phylogenetic non-independence. Both models show a monotonic increase with genetic distance.

**FIGURE 4.**
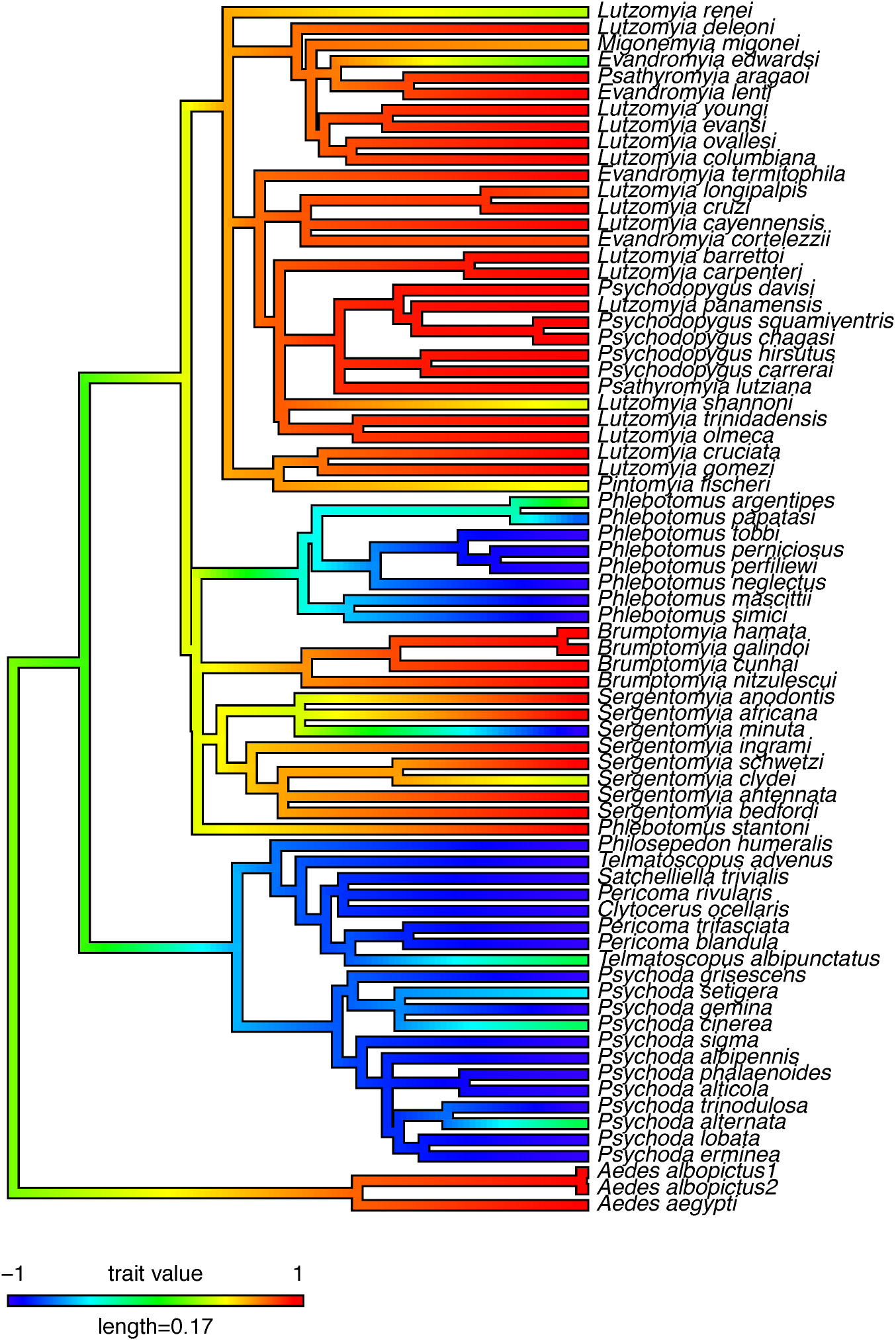
Ancestral reconstruction of TI across the Psychodidae tree shows few transitions in tropicality index. The color of each branch shows the inferred TI values for each node and branch. All ancestral reconstructions used *anc.ML* (*phytools*, (Revell 2012)). Similar trees for PC1 and mean latitude are shown in Figure S4.

#### Models of trait evolution

We fit seven different models of trait evolution to determine which fit best the evolution of climatic niche in the Psychodidae family. These models range from no phylogenetic signal (i.e., white-noise) to punctuated changes of trait evolution associated with speciation events (i.e., *kappa*). Table 3 shows the fit and the parameters inferred for each of the seven models for TI; Tables S4 and S5 show the parameters for PC1 and mean latitude, respectively. Consistent with the results from the summary indices, we find that models with a phylogenetic signal fit better than the only model with no phylogenetic signal for all three proxies of geographic range. When we restricted the comparisons to the white-noise, OU, and Brownian models, the latter model had the lowest AIC for the three proxies of climate niche (Tables 3, S7 and S8). These results suggest evidence for niche conservatism (Wiens et al. 2010). Nonetheless, when we included four additional models, the three proxies differed in the model that best fit their mode of evolution. For PC1, the best fitting model was still a Brownian motion model of trait evolution (*BM* model AIC_BM-PC1_ = 153.618). The best fitting model for mean latitude and TI was the early burst model (*EB* model AIC_EB-Latitude_ = 606.419; *EB* model AIC_EB-TI_ = 99.379) where climatic habitat changes are consistent with the occupation of a variety of climatic niches early in the divergence of the family and a decline of trait evolution as diversification of the family proceeded. These results indicate that even though the evolution of climate niche in Psychodidae has evidence of niche conservatism, macroevolutionary models of trait evolution also suggest the existence of transitions in the phylogenetic history of the group.

**TABLE 3.**
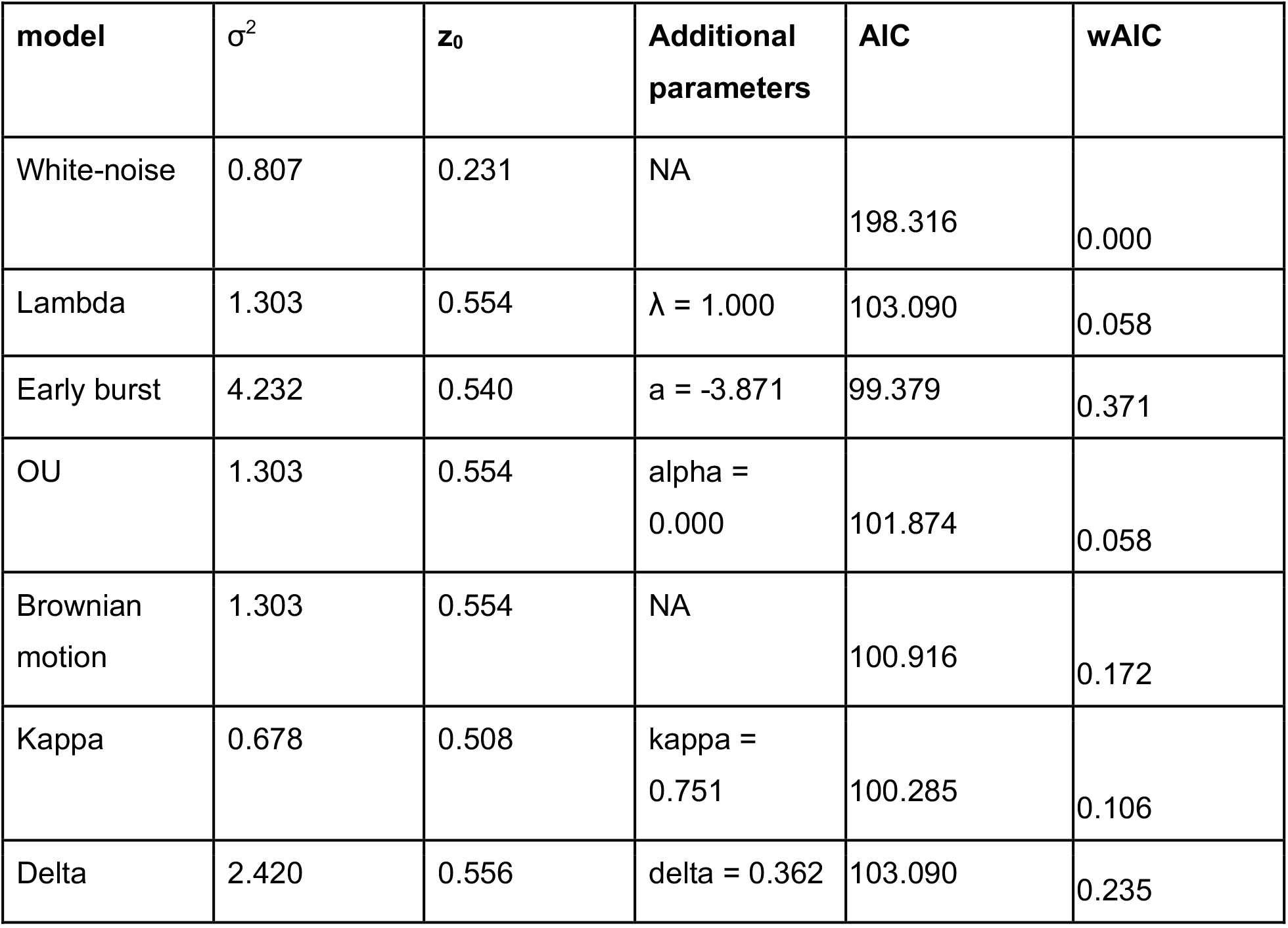
Trait-evolution models suggest that TI in Psychodidae evolves according to an Early-Burst model. σ^2^ is the average amount of change expected in each time step. z0 is the trait value at the root of the tree. Similar analyses and results for PC1 and mean latitude are shown in Table S4 and S5.

#### Ancestral trait reconstruction and transition rates

Figure 4 shows the extant tropicality indexes in the Psychodidae family (marked by color) and the inferred states along the phylogenetic tree. Figure S4 shows similar trees for PC1 and mean latitude. Our best estimate is that the ancestor of the Psychodidae family had a semitropical to tropical distribution (inferred state at the root for tropicality index under a BM model of trait evolution, *Z*_0_ = 0.54; *Z*_0_ values with other models are listed in Table 3). Extant subtropical and temperate species, such as most moth-flies, have a climatic niche that appears to be derived in the family. The clade encompassing vector genera also has an inferred semitropical origin (inferred TI value = 0.39). Within *Phlebotomus*, ancestral character reconstruction suggests that the ancestor of the genus had a semitemperate distribution (inferred TI value = −0.26), suggesting that the colonization of temperate habitats is derived from a semitemperate or semitropical ancestor.

In general, we found that evolutionary transitions to new latitudinal ranges were rare (Figure 5). 92% of transitions from tropical ancestors and 91% of transitions from temperate ancestors resulted in descendant nodes or tips remaining in the same latitudinal zone. Further, all of the remaining 8% of transitions from tropical nodes and 9% of transitions from temperate nodes were to semitropical and semitemperate zones, respectively. By contrast, transitions from the more intermediately defined semitropical and semitemperate zones to adjacent zones were more common and only 33% and 38%, respectively, of descendants from these nodes shared their ancestors’ states.

**FIGURE 5.**
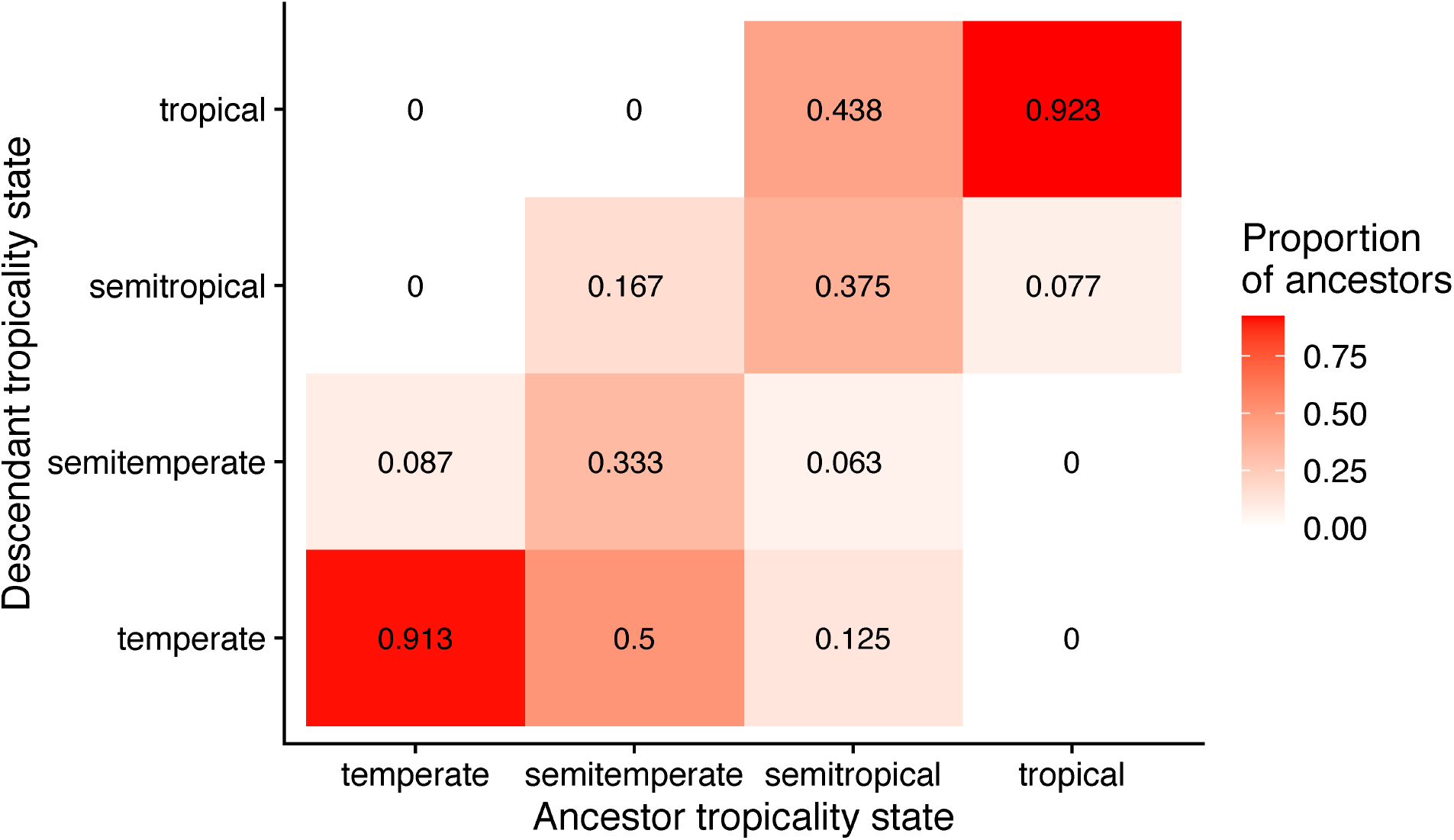
Transitions between latitudinal states, as defined by the tropicality index (TI), are rare in the Psychodidae family. The color of a cell reflects the likelihood of a transition rate. Values in the diagonal represent the likelihood of staying in the same latitudinal band.

## DISCUSSION

In this report, we used georeferenced collections and studied the evolution of climate niche in the Psychodidae family which includes *Leishmania-vector* species. Even though the family distribution spans the tropics to the Arctic circle, our results suggest that different genera within the Psychodidae differ in their climatic niche. Within the family, most species have either a tropical or temperate distribution but rarely span the full hemisphere. Moreover, the phylogenetic distribution of climatic niche components suggests that climate niche has undergone few transitions in Psychodidae. As they have diversified, tropical species have mostly produced tropical species, and temperate species have given rise to more temperate species.

Studies on the evolution of niche divergence as genetic divergence proceeds are rare. In California Jewelflowers, habitat isolation accumulates quickly and remains high (Christie and Strauss 2018) and acts as a barrier against hybridization. This result is qualitatively similar to our findings in *Lutzomyia*, as climatic niche becomes more differentiated as divergence increases. Nonetheless, the scale of the ranges of divergence time differs between the two studies. The study of climatic differentiation in Jewelflowers aimed to understand how barriers to hybridization accumulate between potential interbreeding species, and our goal was to assess the extent of climate differentiation across the whole family, regardless of whether they hybridize or not. Systematic assessments of the magnitude of climatic differentiation across taxa, including those considering the potential effects of climate niche differentiation as a barrier to gene flow via hybridization, are sorely needed to measure the rate of evolution of niche differentiation across the tree of life.

The work presented here has several limitations. Our inference on phylogenetic relationships is largely consistent with previous efforts but should be taken as preliminary. Only a more comprehensive sampling of the variation of the genomes in these dipterans will reveal the true phylogenetic relationships between species. This lack of a fully resolved phylogeny also might affect our results. mtDNA provides low resolution in instances where there has been introgression (e.g., (McVay et al. 2017)). A genome-wide phylogenetic tree that reconstructs the species tree without the limitations of a single gene genealogy is sorely needed. A logical next step in the research of thermal niche differentiation will be to assess whether different species have differences in their realized thermal physiology (reviewed in (Angilletta Jr et al. 2002; Bennett et al. 2019)), in their thermal preference (e.g., (Matute et al. 2009; Cooper et al. 2018)), or in both. Also, the number of transitions from a tropical or temperate node is higher (78 and 46 respectively) than from a semitemperate or semitropical ancestor (6 and 16 respectively) and we thus have more power to detect differences in transition rates between the former categories. Integrating physiological and performance-based traits with analyses of climatic niche evolution can provide a window to understanding differences between physiological and realized niches, ultimately revealing the ecological implications of climatic divergence (Gunderson et al. 2018).

Despite these caveats, our finding of climate niche evolution in the Psychodidae family opens the possibility of new research avenues. First, incorporating a climatic dimension to the study of the evolution of vectors can inform to what extent climate plays a role in the coevolution of parasites and vectors. In the specific case of *Lutzomyia* and *Phlebotomus*, these studies will reveal whether there is an association between carrying *Leishmania* and a tropical climate niche. Second, studies that address the limits of climate niche will also inform which vectors are most likely to move across climatic zones as climate change changes the thermal characteristics of the planet. Finally, comparing the rates of transition between different latitudinal categories can inform whether different taxa show different rates of conservatism. Only one other study has calculated the rates of transition between latitudinal zones (angiosperms; (Kerkhoff et al. 2014)) but the comparison is still informative. The rates of transitions we observed among tropicality values obtained for Psychodidae and those observed for angiosperms are similar and both reveal strong niche conservatism.

As climatic shifts occur globally, changes in environmental conditions will lead to new species distributions (Hitch and Leberg 2007; Rosenberg et al. 2019) or, in the extreme, extinction (Møller et al. 2008). Of particular importance for human health are potential changes in disease vector ranges and abundances, which depend on the extent to which disease vectors exhibit niche conservatism, a phenomenon still poorly examined. Cunze and colleagues (Cunze et al. 2018) suggested niche conservatism in two species of *Aedes* mosquitoes, the vectors of dengue, because none of the two species has filled the entirety of the ecological niche in areas where they have recently invaded. While this evidence shows that *Aedes* have the potential to expand their range in the near future, it does not inform about the extent of conservatism in the genus. Pairwise comparisons of the ecological niche of six pairs of triatomid bugs, vectors of Chagas disease (Ibarra-Cerdeña et al. 2014) suggest that pairs of related species are more similar in their niche than pairs of distantly-related species. While these tests have revealed that niche conservatism might exist for the few species pairs that have been surveyed, the evidence for niche conservatism at the phylogenetic level is still scant. Studies on the climate evolution of species, and in particular of vectors, are important, because species in clades with phylogenetically conserved climatic niches are more likely to shift their geographic distributions in response to changing climate (Tingley et al. 2009, La Sorte and Jetz 2012, Martinez-Meyer and Peterson 2006, Oliveira et al. 2017), raising the possibility of poleward shifts of many vector species that are currently confined to the tropics. Some species of *Lutzomyia*, for example, have expanded their range northward (Comer et al. 1994; Reeves et al. 2008; Minter et al. 2009; Florin et al. 2011; Florin and Rebollar-Téllez 2013) which could potentially expand the endemicity of leishmaniasis (Grosjean et al. 2003; Rosypal et al. 2003, 2005; Schaut et al. 2015).

Understanding the underlying causes of species distribution can inform how species respond to climate change. This is of particular importance to understand how vectors of disease will be distributed around the globe as global warming progresses. Modelling of potential occurrence has revealed that increasing temperatures might increase the potential range of a handful of species (Andrade-Filho et al. 2017; da Costa et al. 2018). Our results suggest that most Psychodidae vector species show a tropical distribution and that an assessment of the potential range expansions given multiple temperature change scenarios might be useful to monitor how vectors expand their niche along latitude and altitude.

## Acknowledgements

We thank J. Coughlan, A. Dagilis, the Matute lab, and two anonymous reviewers for constructive feedback on the manuscript. This work was supported by the National Science Foundation (Dimensions of Biodiversity award 1737752 to D.R.M.). The funders had no role in any aspect of study design, data collection and analysis, or decisions with respect to publication. We have no conflict of interests.

