## Supplementary material for "Climatic niche conservatism in a clade of disease vectors (Diptera: Phlebotominae)": Tables S1-S8, Figures S1-S4

**TABLE S1. Data accessibility.** DOIs for the geographical occurrences of each species included in this study.

| **Species** | **GBIF DOI** |
|---|---|
| *Evandromyia monstruosa* | DOI10.15468/dl.9wru6b |
| *Evandromyia walkeri* | DOI10.15468/dl.ajjbq7 |
| *Evandromyia infraspinosa* | DOI10.15468/dl.hjt3x9 |
| *Psychodopygus amazonensis* | DOI10.15468/dl.m6pvwx |
| *Psychodopygus claustrei* | DOI10.15468/dl.tcrtas |
| *Psychodopygus chagasi* | DOI10.15468/dl.f7yxj6 |
| *Lutzomyia pia* | DOI10.15468/dl.6k7n72 |
| *Lutzomyia hartmanni* | DOI10.15468/dl.8cykdb |
| *Lutzomyia anthophora* | DOI10.15468/dl.twync2 |
| *Lutzomyia trapidoi* | DOI10.15468/dl.dfu35p |
| *Lutzomyia verrucarum* | DOI10.15468/dl.hjuk4v |
| *Lutzomyia whitmani* | DOI10.15468/dl.6zr2tu |
| *Trichomyia fusca* | DOI10.15468/dl.v44uqe |
| *Trichomyia saurotis* | DOI10.15468/dl.jrc4mq |
| *Trichomyia acanthostylis* | DOI10.15468/dl.ctjxrd |
| *Trichomyia xaniostylis* | DOI10.15468/dl.ahp69x |
| *Trichomyia dolichakis* | DOI10.15468/dl.gqettr |
| *Trichomyia brevitarsa* | DOI10.15468/dl.e6y4pv |
| *Trichomyia nuda* | DOI10.15468/dl.jfdn6w |
| *Trichomyia urbica* | DOI10.15468/dl.jgbwws |
| *Philosepedon opposita* | DOI10.15468/dl.pzpc2d |

| | |
|---|---|
| *Philosepedon interdicta* | DOI10.15468/dl.zemg6a |
| *Philosepedon balkanicus* | DOI10.15468/dl.fxqw6n |
| *Warileya rotundipennis* | DOI10.15468/dl.xagvet |
| *Clytocerus dalii* | DOI10.15468/dl.rsr28e |
| *Clytocerus rivosus* | DOI10.15468/dl.eaau49 |
| *Clytocerus tetracorniculatus* | DOI10.15468/dl.yqd9m7 |
| *Clytocerus splendidus* | DOI10.15468/dl.5bkvcs |
| *Clytocerus ocellaris* | DOI10.15468/dl.axbdtw |
| *Satchelliella crispi* | DOI10.15468/dl.erqm4c |
| *Satchelliella gracilis* | DOI10.15468/dl.vsv4pv |
| *Satchelliella palustris* | DOI10.15468/dl.nmwyc8 |
| *Satchelliella nubila* | DOI10.15468/dl.szvhjf |
| *Satchelliella extricata* | DOI10.15468/dl.dmyhmv |
| *Satchelliella pilularia* | DOI10.15468/dl.ds3nwg |
| *Satchelliella ussurica* | DOI10.15468/dl.anqquv |
| *Satchelliella compta* | DOI10.15468/dl.qtufbf |
| *Satchelliella stammeri* | DOI10.15468/dl.qghju5 |
| *Satchelliella borealis* | DOI10.15468/dl.s9b84m |
| *Satchelliella mutua* | DOI10.15468/dl.m8up6a |
| *Satchelliella trivialis* | DOI10.15468/dl.92psqg |
| *Psychodopygus paraensis* | DOI10.15468/dl.uz74t6 |
| *Psychodopygus paraensis* | DOI10.15468/dl.ehrpv5 |
| *Psychodopygus chagasi* | DOI10.15468/dl.7c8mmf |
| *Psychodopygus carrerai* | DOI10.15468/dl.xthkcv |
| *Psychodopygus squamiventris* | DOI10.15468/dl.q24zhy |
| *Psychodopygus hirsutus* | DOI10.15468/dl.8mmtpa |
| *Psychodopygus davisi* | DOI10.15468/dl.h3f3ee |
| *Psychodopygus davisi* | DOI10.15468/dl.2zgk7d |

| | |
|---|---|
| *Psathyromyia inflata* | DOI10.15468/dl.cqqwrr |
| *Psathyromyia inflata* | DOI10.15468/dl.u8vtqy |
| *Psathyromyia campograndensis* | DOI10.15468/dl.5utxqt |
| *Psathyromyia barrettoi* | DOI10.15468/dl.m6dvxy |
| *Psathyromyia aragaoi* | DOI10.15468/dl.2dmxj7 |
| *Brumptomyia travassosi* | DOI10.15468/dl.3qkkvw |
| *Brumptomyia beaupertuyi* | DOI10.15468/dl.nunnae |
| *Brumptomyia guimaraesi* | DOI10.15468/dl.xpsr3g |
| *Brumptomyia ortizi* | DOI10.15468/dl.f4tadw |
| *Brumptomyia troglodytes* | DOI10.15468/dl.38yqv3 |
| *Brumptomyia brumpti* | DOI10.15468/dl.u6rguk |
| *Brumptomyia pintoi* | DOI10.15468/dl.fnapj6 |
| *Brumptomyia pentacantha* | DOI10.15468/dl.yjj6nq |
| *Brumptomyia nitzulescui* | DOI10.15468/dl.mnfjyc |
| *Brumptomyia cunhai* | DOI10.15468/dl.e3xunm |
| *Brumptomyia figueiredoi* | DOI10.15468/dl.zjwjxv |
| *Brumptomyia avellari* | DOI10.15468/dl.buu22m |
| *Brumptomyia hamata* | DOI10.15468/dl.nnqgzr |
| *Migonemyia vaniae* | DOI10.15468/dl.62cjkg |
| *Migonemyia migonei* | DOI10.15468/dl.awcvew |
| *Phlebotomus longiductus* | DOI10.15468/dl.c9bjd6 |
| *Phlebotomus transcaucasicus* | DOI10.15468/dl.e2ggwp |
| *Phlebotomus aculeatus* | DOI10.15468/dl.jpn86b |
| *Phlebotomus bergeroti* | DOI10.15468/dl.3ygk4p |
| *Phlebotomus langeroni* | DOI10.15468/dl.wmt24m |
| *Phlebotomus kandelakii* | DOI10.15468/dl.539edt |
| *Phlebotomus mascittii* | DOI10.15468/dl.sy25g8 |
| *Phlebotomus alexandri* | DOI10.15468/dl.x3e4jf |

| | |
|---|---|
| *Phlebotomus ariasi* | DOI10.15468/dl.hbx8cb |
| *Phlebotomus mascomai* | DOI10.15468/dl.hgtcab<br>DOI10.15468/dl.aac8qh |
| *Pericoma barbata* | DOI10.15468/dl.8sdw35 |
| *Pericoma gourlayi* | DOI10.15468/dl.s9963m |
| *Pericoma bifalcata* | DOI10.15468/dl.j468zm |
| *Pericoma diffusa* | DOI10.15468/dl.4wnz8b |
| *Pericoma tenerifensis* | DOI10.15468/dl.9xscf3 |
| *Pericoma isabellae* | DOI10.15468/dl.rubvt7 |
| *Pericoma albitarsis* | DOI10.15468/dl.7e57hx |
| *Pericoma spiralifera* | DOI10.15468/dl.ejm86s |
| *Pericoma marginalis* | DOI10.15468/dl.9t4p9s |
| *Pericoma soleata* | DOI10.15468/dl.ag5unb |
| *Pericoma multimaculata* | DOI10.15468/dl.ft24mx |
| *Pericoma bipunctata* | DOI10.15468/dl.fvmguq |
| *Pericoma zumbadoi* | DOI10.15468/dl.fpq5fq |
| *Pericoma fallax* | DOI10.15468/dl.8nppez |
| *Pericoma nigricauda* | DOI10.15468/dl.q5bfpt |
| *Pericoma californica* | DOI10.15468/dl.bve68w |
| *Pericoma hespenheidei* | DOI10.15468/dl.g9sn7g |
| *Pericoma exquisita* | DOI10.15468/dl.z29myf |
| *Pericoma trifasciata* | DOI10.15468/dl.p7g85h |
| *Pericoma pseudoexquisita* | DOI10.15468/dl.m5b89n |
| *Pericoma albitarsis* | DOI10.15468/dl.m92zt8 |
| *Pericoma americana* | DOI10.15468/dl.2ddue7 |
| *Pericoma nielseni* | DOI10.15468/dl.h9xawy |
| *Pericoma blandula* | DOI10.15468/dl.bsa8gx |
| *Pericoma rivularis* | DOI10.15468/dl.kkzykr |

| | |
|---|---|
| *Evandromyia termitophila* | DOI10.15468/dl.3zr56n |
| *Evandromyia cortelezzii* | DOI10.15468/dl.jvfpys |
| *Evandromyia lenti* | DOI10.15468/dl.47q697 |
| *Telmatoscopus labeculosus* | DOI10.15468/dl.9vfwxf |
| *Telmatoscopus labeculosus* | DOI10.15468/dl.9vfwxf |
| *Telmatoscopus britteni* | DOI10.15468/dl.jcbbua |
| *Telmatoscopus laurenci* | DOI10.15468/dl.bf2v29 |
| *Telmatoscopus vaillanti* | DOI10.15468/dl.xy5mxx |
| *Telmatoscopus morulus* | DOI10.15468/dl.j579ey |
| *Telmatoscopus ambiguus* | DOI10.15468/dl.z2a5ku |
| *Telmatoscopus niger* | DOI10.15468/dl.vtaq2g |
| *Telmatoscopus basalis* | DOI10.15468/dl.yjv6z2 |
| *Telmatoscopus similis* | DOI10.15468/dl.dwfd2q |
| *Telmatoscopus advenus* | DOI10.15468/dl.hzss4b |
| *Telmatoscopus rothschildi* | DOI10.15468/dl.j79kks |
| *Telmatoscopus furcatus* | DOI10.15468/dl.z977tj |
| *Telmatoscopus superbus* | DOI10.15468/dl.q7dnpq |
| *Telmatoscopus albipunctata* | DOI10.15468/dl.pdzbgw |
| *Pintomyia paleotrichia* | DOI10.15468/dl.r26wcu |
| *Pintomyia falcaorum* | DOI10.15468/dl.b54e3q |
| *Pintomyia fischeri* | DOI10.15468/dl.s5cu3u |
| *Micropygomyia trinidadensis* | DOI10.15468/dl.mrj4ed |
| *Micropygomyia peresi* | DOI10.15468/dl.vuv3cr |
| *Micropygomyia micropyga* | DOI10.15468/dl.uz5zq6 |
| *Sergentomyia suberecta* | DOI10.15468/dl.t2fr3d |
| *Sergentomyia adleri* | DOI10.15468/dl.y22wxg |
| *Sergentomyia bailyi* | DOI10.15468/dl.a77xtg |
| *Sergentomyia affinis* | DOI10.15468/dl.3xzwfd |

| | |
|---|---|
| *Sergentomyia mangana* | DOI10.15468/dl.2qyata |
| *Sergentomyia decipiens* | DOI10.15468/dl.ytbwuh |
| *Sergentomyia garnhami* | DOI10.15468/dl.tuxevz |
| *Sergentomyia africana* | DOI10.15468/dl.rdg7ub |
| *Sergentomyia minuta* | DOI10.15468/dl.cwdasp |
| *Sergentomyia clydei* | DOI10.15468/dl.5gb2m5 |
| *Sergentomyia ingrami* | DOI10.15468/dl.judd87 |
| *Sergentomyia schwetzi* | DOI10.15468/dl.cyk4b4 |
| *Sergentomyia antennata* | DOI10.15468/dl.d5hwh8 |
| *Sergentomyia bedfordi* | DOI10.15468/dl.pmpse2 |
| *Psychoda pusilla* | DOI10.15468/dl.3kgwju |
| *Psychoda alticola* | DOI10.15468/dl.27x7qr |
| *Psychoda crassipennis* | DOI10.15468/dl.edur2t |
| *Psychoda cultella* | DOI10.15468/dl.r4uy2s |
| *Psychoda pitilla* | DOI10.15468/dl.jnbp7g |
| *Psychoda quadrifilis* | DOI10.15468/dl.x2666s |
| *Psychoda sigma* | DOI10.15468/dl.bw9fap |
| *Psychoda fasciata* | DOI10.15468/dl.rsz8m4 |
| *Psychoda erminea* | DOI10.15468/dl.rvxbsx |
| *Psychoda entolopha* | DOI10.15468/dl.w9r7fq |
| *Psychoda itoco* | DOI10.15468/dl.ff2zhr |
| *Psychoda surcoufi* | DOI10.15468/dl.rnekt2 |
| *Psychoda savaiiensis* | DOI10.15468/dl.wfzhbj |
| *Psychoda minuta* | DOI10.15468/dl.3cw3fn |
| *Psychoda setigera* | DOI10.15468/dl.sdzbeg |
| *Psychoda alternicula* | DOI10.15468/dl.wqyyxn |
| *Psychoda tothastica* | DOI10.15468/dl.wq3hkg |
| *Psychoda lobata* | DOI10.15468/dl.2hueby |

| | |
|---|---|
| *Psychoda grisescens* | DOI10.15468/dl.su6zqu |
| *Psychoda gemina* | DOI10.15468/dl.226zzv |
| *Psychoda satchelli* | DOI10.15468/dl.qag8fr |
| *Psychoda cinerea* | DOI10.15468/dl.ygupmr |
| *Psychoda albipennis* | DOI10.15468/dl.8939rr |
| *Psychoda alternata* | DOI10.15468/dl.eav3v6 |
| *Psychoda trinodulosa* | DOI10.15468/dl.ttb88q |
| *Psychoda phalaenoides* | DOI10.15468/dl.c398yp |
| *Phlebotomus rodhaini* | DOI10.15468/dl.ckgn5j |
| *Phlebotomus perfiliewi* | DOI10.15468/dl.mq2vas |
| *Phlebotomus argentipes* | DOI10.15468/dl.mgp7eq |
| *Phlebotomus stantoni* | DOI10.15468/dl.e4h4sp |
| *Phlebotomus guggisbergi* | DOI10.15468/dl.eyx6xb |
| *Phlebotomus longicuspis* | DOI10.15468/dl.xsrq7h |
| *Phlebotomus martini* | DOI10.15468/dl.xpxt99 |
| *Phlebotomus celiae* | DOI10.15468/dl.tsfn36 |
| *Phlebotomus saevus* | DOI10.15468/dl.ky8e9n |
| *Phlebotomus duboscqi* | DOI10.15468/dl.3hdst3 |
| *Phlebotomus elgonensis* | DOI10.15468/dl.za4n3m |
| *Phlebotomus tobbi* | DOI10.15468/dl.wcee8m |
| *Phlebotomus simici* | DOI10.15468/dl.9k9dvy |
| *Phlebotomus sergenti* | DOI10.15468/dl.y6y6wp |
| *Phlebotomus neglectus* | DOI10.15468/dl.xsu4gx |
| *Phlebotomus pedifer* | DOI10.15468/dl.9896dv |
| *Phlebotomus perniciosus* | DOI10.15468/dl.bufazu |
| *Phlebotomus papatasi* | DOI10.15468/dl.7fswbg |
| *Phlebotomus similis* | DOI10.15468/dl.8e4trb |
| *Lutzomyia cayennensis* | DOI10.15468/dl.83g8kp |

| | |
|---|---|
| *Lutzomyia chiapanensis* | DOI10.15468/dl.hra7bp |
| *Lutzomyia edwardsi* | DOI10.15468/dl.e55ytw |
| *Lutzomyia ubiquitalis* | DOI10.15468/dl.w9t49q |
| *Lutzomyia lainsoni* | DOI10.15468/dl.vmtbx3 |
| *Lutzomyia ininii* | DOI10.15468/dl.vvz4x2 |
| *Lutzomyia yencanensis* | DOI10.15468/dl.5peb94 |
| *Lutzomyia ischyracantha* | DOI10.15468/dl.t6jf8p |
| *Lutzomyia damascenoi* | DOI10.15468/dl.sy4ggg |
| *Lutzomyia caprina* | DOI10.15468/dl.cjt4wc |
| *Lutzomyia geniculata* | DOI10.15468/dl.488r8u |
| *Lutzomyia carpenteri* | DOI10.15468/dl.h8j6uy |
| *Lutzomyia cruzi* | DOI10.15468/dl.etgc7n |
| *Lutzomyia lutziana* | DOI10.15468/dl.zctcnj |
| *Lutzomyia goiana* | DOI10.15468/dl.qrrfju |
| *Lutzomyia ovallesi* | DOI10.15468/dl.n4j9nx |
| *Lutzomyia migonei* | DOI10.15468/dl.xrj6nf |
| *Lutzomyia ischnacantha* | DOI10.15468/dl.624xsb |
| *Lutzomyia cavernicola* | DOI10.15468/dl.cms9d5 |
| *Lutzomyia monticola* | DOI10.15468/dl.cf7q2m |
| *Lutzomyia renei* | DOI10.15468/dl.3s5esu |
| *Lutzomyia dendrophyla* | DOI10.15468/dl.kr2r44 |
| *Lutzomyia deleoni* | DOI10.15468/dl.jbv4a5 |
| *Lutzomyia cruciata* | DOI10.15468/dl.hajavv |
| *Lutzomyia barrettoi* | DOI10.15468/dl.gy6a4j |
| *Lutzomyia ayrozai* | DOI10.15468/dl.sh6nsr |
| *Lutzomyia yuilli* | DOI10.15468/dl.b4u42h |
| *Lutzomyia micropyga* | DOI10.15468/dl.gxm4yp |
| *Lutzomyia shannoni* | DOI10.15468/dl.kru9ks |

| | |
|---|---|
| *Lutzomyia panamensis* | DOI10.15468/dl.svwgzn |
| *Lutzomyia atroclavata* | DOI10.15468/dl.n6uud4 |
| *Lutzomyia venezuelensis* | DOI10.15468/dl.jverbw |
| *Lutzomyia trinidadensis* | DOI10.15468/dl.keu2s9 |
| *Lutzomyia cayennensis* | DOI10.15468/dl.xhzbrh |
| *Lutzomyia rangeliana* | DOI10.15468/dl.yy7knb |
| *Lutzomyia gomezi* | DOI10.15468/dl.wyjnqu |
| *Lutzomyia youngi* | DOI10.15468/dl.gezw5h |
| *Lutzomyia columbiana* | DOI10.15468/dl.39cbaa |
| *Lutzomyia olmeca* | DOI10.15468/dl.pd69c5 |
| *Lutzomyia townsendi* | DOI10.15468/dl.jet5h2 |
| *Lutzomyia evansi* | DOI10.15468/dl.cmw6fz |
| *Lutzomyia longipalpis* | DOI10.15468/dl.rssvqb |

**TABLE S2. Genebank accession numbers used to generate a phylogenetic tree for the Psychodidae family.**

| **Genus** | **Species** | **Genebank number** |
|---|---|---|
| *Brumptomyia* | *Brumptomyia guimaraesi* | KC921225.1 |
| *Brumptomyia* | *Brumptomyia mesai* | MK851242.1 |
| *Brumptomyia* | *Brumptomyia hamata* | KR907866.1 |
| *Brumptomyia* | *Brumptomyia avellari* | MG234309.1 |
| *Brumptomyia* | *Brumptomyia beaupertuyi* | KC921222.1 |
| *Brumptomyia* | *Brumptomyia galindoi* | GU001735.1 |
| *Brumptomyia* | *Brumptomyia ortizi* | KP112523.1 |
| *Brumptomyia* | *Brumptomyia nitzulescui* | KP112520.1 |
| *Brumptomyia* | *Brumptomyia cunhai* | KP112505.1 |

| | | |
|---|---|---|
| *Clytocerus* | *Clytocerus ocellaris* | MF966155.1 |
| *Lutzomyia* | *Lutzomyia carpenteri* | KC921236.1 |
| *Lutzomyia* | *Lutzomyia cayennensis* | GU909475.1 |
| *Lutzomyia* | *Lutzomyia columbiana* | KC921246.1 |
| *Lutzomyia* | *Lutzomyia cruciata* | MK851248.1 |
| *Lutzomyia* | *Lutzomyia cruzi* | KP112576.1 |
| *Lutzomyia* | *Lutzomyia deleoni* | MK851253.1 |
| *Lutzomyia* | *Lutzomyia evansi* | JN845538.1 |
| *Lutzomyia* | *Lutzomyia gomezi* | KC921254.1 |
| *Lutzomyia* | *Lutzomyia guyanensis* | GU001736.1 |
| *Lutzomyia* | *Lutzomyia inni* | KX356040.1 |
| *Lutzomyia* | *Lutzomyia longipalpis* | MK851273.1 |
| *Lutzomyia* | *Lutzomyia lutziana* | KP112939.1 |
| *Lutzomyia* | *Lutzomyia micropyga* | GU909466.1 |
| *Lutzomyia* | *Lutzomyia migonei* | GU909509.1 |
| *Lutzomyia* | *Lutzomyia olmeca* | GU001743.1 |
| *Lutzomyia* | *Lutzomyia ovallesi* | MN257603.1 |
| *Lutzomyia* | *Lutzomyia panamensis* | GU909463.1 |
| *Lutzomyia* | *Lutzomyia rangeliana* | GU909494.1 |
| *Lutzomyia* | *Lutzomyia renei* | KP112605.1 |
| *Lutzomyia* | *Lutzomyia shannoni* | KC921296.1 |
| *Lutzomyia* | *Lutzomyia trinidadensis* | KC921309.1 |
| *Lutzomyia* | *Lutzomyia venezuelensis* | GU909484.1 |
| *Lutzomyia* | *Lutzomyia youngi* | KC921327.1 |
| *Lutzomyia* | *Lutzomyia yuilli* | KC921326.1 |
| *Lutzomyia* | *Lutzomyia whitmani* | MG234412.1 |
| *Lutzomyia* | *Lutzomyia verrucarum* | AB984470.1 |

| | | |
|---|---|---|
| *Lutzomyia* | *Lutzomyia trapidoi* | KC921305.1 |
| *Lutzomyia* | *Lutzomyia anthophora* | MK952678.1 |
| *Lutzomyia* | *Lutzomyia hartmanni* | KC921258.1 |
| *Lutzomyia* | *Lutzomyia atroclavata* | GU909482.1 |
| *Lutzomyia* | *Lutzomyia pia* | KC921280.1 |
| *Lutzomyia* | *Psychodopygus ayrozai* | KP112967.1 |
| *Trichomyia* | *Trichomyia sp. P44* | MH042541.1 |
| *Trichomyia* | *Trichomyia ituberensis* | MH042540.1 |
| *Trichomyia* | *Trichomyia pseudoannae* | MH042536.1 |
| *Trichomyia* | *Trichomyia cerdosa* | MH042537.1 |
| *Philosepedon* | *Philosepedon humeralis* | MH917737.1 |
| *Philosepedon* | *Philosepedon sp. 1* | JQ349584.1 |
| *Warileya* | *Warileya rotundipennis* | KC921329.1 |
| *Warileya* | *Warileya phlebotomanica* | AB761141.1 |
| *Warileya* | *Warileya euniceae* | AB984520.1 |
| *Satcheliella* | *Satchelliella trivialis* | MF966157.1 |
| *Psychodopygus* | *Psychodopygus carrerai* | MG029462.1 |
| *Psychodopygus* | *Psychodopygus panamensis* | KC921266.1 |
| *Psychodopygus* | *Psychodopygus hirsutus* | KP112999.1 |
| *Psychodopygus* | *Psychodopygus squamiventris* | MH281929.1 |
| *Psychodopygus* | *Psychodopygus chagasi* | MH281900.1 |
| *Psychodopygus* | *Psychodopygus davisi* | MG234425.1 |
| *Psathyromyia* | *Psathyromyia aragaoi* | KP763471.1 |
| *Psathyromyia* | *Lutzomyia barrettoi* | KC921227.1 |
| *Migomemyia* | *Migonemyia migonei* | MT430890.1 |
| *Phlebotomus* | *Phlebotomus longiductus* | KF137552.1 |
| *Phlebotomus* | *Phlebotomus transcaucasicus* | KY564181.1 |

| | | |
|---|---|---|
| *Phlebotomus* | *Phlebotomus aculeatus* | MK169222.1 |
| *Phlebotomus* | *Phlebotomus bergeroti* | KY883629.1 |
| *Phlebotomus* | *Phlebotomus kandelakii* | KY564184.1 |
| *Phlebotomus* | *Phlebotomus mascittii* | MN003381.1 |
| *Phlebotomus* | *Phlebotomus alexandri* | MN086432.1 |
| *Phlebotomus* | *Phlebotomus ariasi* | MH559443.1 |
| *Phlebotomus* | *Phlebotomus mascomai* | FJ493548.1 |
| *Phlebotomus* | *Phlebotomus rodhaini* | JX105042.1 |
| *Phlebotomus* | *Phlebotomus perfiliewi* | MG948469.1 |
| *Phlebotomus* | *Phlebotomus argentipes* | KY883622.1 |
| *Phlebotomus* | *Phlebotomus stantoni* | MF966710.1 |
| *Phlebotomus* | *Phlebotomus guggisbergi* | MK169221.1 |
| *Phlebotomus* | *Phlebotomus longicuspis* | KJ481168.1 |
| *Phlebotomus* | *Phlebotomus martini* | JX105040 |
| *Phlebotomus* | *Phlebotomus celiae* | JX105041 |
| *Phlebotomus* | *Phlebotomus saevus* | KF483673.1 |
| *Phlebotomus* | *Phlebotomus duboscqi* | KY911406.1 |
| *Phlebotomus* | *Phlebotomus tobbi* | MN086651.1 |
| *Phlebotomus* | *Phlebotomus simici* | MN086717.1 |
| *Phlebotomus* | *Phlebotomus sergenti* | KC755398.1 |
| *Phlebotomus* | *Phlebotomus neglectus* | MN003379.1 |
| *Phlebotomus* | *Phlebotomus perniciosus* | LC090044.1 |
| *Phlebotomus* | *Phlebotomus papatasi* | MH780862.1 |
| *Phlebotomus* | *Phlebotomus similis* | MF968982.1 |
| *Pericoma* | *Pericoma blandula* | KF549514.1 |
| *Pericoma* | *Pericoma rivularis* | KF549497.1 |
| *Pericoma* | *Pericoma trifasciata* | MH592661.1 |

| | | |
|---|---|---|
| *Evandromyia* | *Evandromyia infraspinosa* | MG234326.1 |
| *Evandromyia* | *Evandromyia lenti* | MN181553.1 |
| *Evandromyia* | *Evandromyia edwardsi* | KP112547.1 |
| *Evandromyia* | *Evandromyia termitophila* | MG234348.1 |
| *Evandromyia* | *Evandromyia walkeri* | MN181565.1 |
| *Evandromyia* | *Evandromyia cortelezzii* | MN432911.1 |
| *Telmatoscopus* | *Telmatoscopus advenus* | KF549511.1 |
| *Telmatoscopus* | *Telmatoscopus albipunctatus* | KY924868.1 |
| *Pintomyia* | *Pintomyia fischeri* | KP112811.1 |
| *Pintomyia* | *Pintomyia monticola* | KP112884.1 |
| *Micropygomyia* | *Micropygomyia trinidadensis* | MG234381.1 |
| *Micropygomyia* | *Micropygomyia micropyga* | KR907857.1 |
| *Micropygomyia* | *Micropygomyia peresi* | KP112637.1 |
| *Sengentomyia* | *Sergentomyia minuta* | KP828572.1 |
| *Sengentomyia* | *Sergentomyia schwetzi* | MG913288.1 |
| *Sengentomyia* | *Sergentomyia bailyi i* | KX270830.1 |
| *Sengentomyia* | *Sergentomyia clydei* | MH577119.1 |
| *Sengentomyia* | *Sergentomyia antennata* | MH577082.1 |
| *Sengentomyia* | *Sergentomyia affinis* | MH577066.1 |
| *Sengentomyia* | *Sergentomyia anodontis* | MN850832.1 |
| *Sengentomyia* | *Sergentomyia africana* | MH577100.1 |
| *Sengentomyia* | *Sergentomyia ingrami* | AB759973.1 |
| *Sengentomyia* | *Sergentomyia adleri* | MH577117.1 |
| *Sengentomyia* | *Sergentomyia bedfordi* | KY451795.1 |
| *Psychoda* | *Psychoda trinodulosa* | KT089508.1 |
| *Psychoda* | *Psychoda alternata* | JF872473.1 |
| *Psychoda* | *Psychoda phalaenoides* | MG296209.1 |

| | | |
|---|---|---|
| *Psychoda* | *Psychoda grisescens* | KT089934.1 |
| *Psychoda* | *Psychoda cinerea* | MG298616.1 |
| *Psychoda* | *Psychoda nr. albipennis* | JQ349627.1 |
| *Psychoda* | *Psychoda lobata* | KF549507.1 |
| *Psychoda* | *Psychoda erminea* | KF549506.1 |
| *Psychoda* | *Psychoda sigma* | JN298980.1 |
| *Psychoda* | *Psychoda setigera* | KF549502.1 |
| *Psychoda* | *Psychoda alticola* | KF549500.1 |
| *Psychoda* | *Psychoda gemina* | MF966166.1 |
| *Aedes* | *Aedes albopictus* | KM457564.1 |
| *Aedes* | *Aedes aegyptii* | MK265729.1 |

**TABLE S3**. PC loadings for the Psycodidae family-level PCA.

| | PC1 | PC2 | PC3 | PC4 | PC5 |
|---|---|---|---|---|---|
| bio5_50 | -0.528722 | 0.3140102 | -0.0690605 | 0.2784433 | 0.73454112 |
| bio6_50 | -0.5424083 | 0.1121851 | 0.19645291 | 0.5223093 | -0.6179053 |
| bio12_50 | -0.3788427 | -0.3142702 | 0.7243473 | -0.4704755 | 0.10810298 |
| bio15_50 | -0.47172 | 0.2194634 | -0.5235676 | -0.6289459 | -0.2441722 |
| elev_50 | -0.2453665 | -0.8613258 | -0.3972843 | 0.180948 | 0.08565075 |

**TABLE S4.** Tukey pairwise comparisons for PC1

| **contrast** | **estimate** | **SE** | **df** | **t.ratio** | **p.value** |
|---|---|---|---|---|---|
| *Brumptomyia - Clytocerus* | -3.1205 | 0.529 | 115 | -5.896 | <.0001 |
| *Brumptomyia - Lutzomyia* | 0.4308 | 0.414 | 115 | 1.041 | 0.9936 |

| | | | | | |
|---|---|---|---|---|---|
| *Brumptomyia - Pericoma* | -2.8706 | 0.467 | 115 | -6.15 | <.0001 |
| *Brumptomyia - Philosepedon* | -2.3348 | 0.603 | 115 | -3.875 | 0.0079 |
| *Brumptomyia - Phlebotomus* | -0.3199 | 0.467 | 115 | -0.685 | 0.9998 |
| *Brumptomyia - Psychoda* | -2.356 | 0.427 | 115 | -5.512 | <.0001 |
| *Brumptomyia - Satchelliella* | -3.1885 | 0.467 | 115 | -6.831 | <.0001 |
| *Brumptomyia - Sergentomyia* | 0.6853 | 0.474 | 115 | 1.445 | 0.9339 |
| *Brumptomyia - Telmatoscopus* | -2.6822 | 0.474 | 115 | -5.658 | <.0001 |
| *Brumptomyia - Trichomyia* | -1.5766 | 0.603 | 115 | -2.616 | 0.2528 |
| *Clytocerus - Lutzomyia* | 3.5513 | 0.374 | 115 | 9.49 | <.0001 |
| *Clytocerus - Pericoma* | 0.2499 | 0.432 | 115 | 0.578 | 1 |
| *Clytocerus - Philosepedon* | 0.7857 | 0.576 | 115 | 1.364 | 0.9546 |
| *Clytocerus - Phlebotomus* | 2.8006 | 0.432 | 115 | 6.481 | <.0001 |
| *Clytocerus - Psychoda* | 0.7645 | 0.389 | 115 | 1.964 | 0.6739 |
| *Clytocerus - Satchelliella* | -0.0681 | 0.432 | 115 | -0.157 | 1 |
| *Clytocerus - Sergentomyia* | 3.8058 | 0.44 | 115 | 8.648 | <.0001 |
| *Clytocerus - Telmatoscopus* | 0.4382 | 0.44 | 115 | 0.996 | 0.9955 |
| *Clytocerus - Trichomyia* | 1.5439 | 0.576 | 115 | 2.68 | 0.2223 |
| *Lutzomyia - Pericoma* | -3.3014 | 0.279 | 115 | -11.836 | <.0001 |
| *Lutzomyia - Philosepedon* | -2.7656 | 0.472 | 115 | -5.856 | <.0001 |
| *Lutzomyia - Phlebotomus* | -0.7507 | 0.279 | 115 | -2.691 | 0.217 |
| *Lutzomyia - Psychoda* | -2.7868 | 0.206 | 115 | -13.498 | <.0001 |
| *Lutzomyia - Satchelliella* | -3.6194 | 0.279 | 115 | -12.976 | <.0001 |
| *Lutzomyia - Sergentomyia* | 0.2545 | 0.291 | 115 | 0.874 | 0.9985 |

| | | | | | |
|---|---|---|---|---|---|
| *Lutzomyia - Telmatoscopus* | -3.1131 | 0.291 | 115 | -10.695 | <.0001 |
| *Lutzomyia - Trichomyia* | -2.0074 | 0.472 | 115 | -4.25 | 0.0021 |
| *Pericoma - Philosepedon* | 0.5358 | 0.519 | 115 | 1.032 | 0.9941 |
| *Pericoma - Phlebotomus* | 2.5507 | 0.353 | 115 | 7.229 | <.0001 |
| *Pericoma - Psychoda* | 0.5146 | 0.299 | 115 | 1.722 | 0.8212 |
| *Pericoma - Satchelliella* | -0.318 | 0.353 | 115 | -0.901 | 0.998 |
| *Pericoma - Sergentomyia* | 3.5559 | 0.362 | 115 | 9.809 | <.0001 |
| *Pericoma - Telmatoscopus* | 0.1883 | 0.362 | 115 | 0.52 | 1 |
| *Pericoma - Trichomyia* | 1.294 | 0.519 | 115 | 2.492 | 0.3205 |
| *Philosepedon - Phlebotomus* | 2.0149 | 0.519 | 115 | 3.88 | 0.0078 |
| *Philosepedon - Psychoda* | -0.0212 | 0.484 | 115 | -0.044 | 1 |
| *Philosepedon - Satchelliella* | -0.8538 | 0.519 | 115 | -1.644 | 0.8598 |
| *Philosepedon - Sergentomyia* | 3.0201 | 0.526 | 115 | 5.742 | <.0001 |
| *Philosepedon - Telmatoscopus* | -0.3475 | 0.526 | 115 | -0.661 | 0.9999 |
| *Philosepedon - Trichomyia* | 0.7582 | 0.644 | 115 | 1.177 | 0.9838 |
| *Phlebotomus - Psychoda* | -2.0361 | 0.299 | 115 | -6.813 | <.0001 |
| *Phlebotomus - Satchelliella* | -2.8686 | 0.353 | 115 | -8.13 | <.0001 |
| *Phlebotomus - Sergentomyia* | 1.0052 | 0.362 | 115 | 2.773 | 0.1821 |
| *Phlebotomus - Telmatoscopus* | -2.3623 | 0.362 | 115 | -6.517 | <.0001 |
| *Phlebotomus - Trichomyia* | -1.2567 | 0.519 | 115 | -2.42 | 0.3636 |
| *Psychoda - Satchelliella* | -0.8325 | 0.299 | 115 | -2.786 | 0.1769 |
| *Psychoda - Sergentomyia* | 3.0413 | 0.31 | 115 | 9.804 | <.0001 |
| *Psychoda - Telmatoscopus* | -0.3262 | 0.31 | 115 | -1.052 | 0.9931 |

| | | | | | |
|---|---|---|---|---|---|
| *Psychoda - Trichomyia* | 0.7794 | 0.484 | 115 | 1.609 | 0.8752 |
| *Satchelliella - Sergentomyia* | 3.8738 | 0.362 | 115 | 10.686 | <.0001 |
| *Satchelliella - Telmatoscopu* | s 0.5063 | 0.362 | 115 | 1.397 | 0.9469 |
| *Satchelliella - Trichomyia* | 1.612 | 0.519 | 115 | 3.104 | 0.0818 |
| *Sergentomyia - Telmatoscopus* | -3.3675 | 0.372 | 115 | -9.055 | <.0001 |
| *Sergentomyia - Trichomyia* | -2.2619 | 0.526 | 115 | -4.3 | 0.0017 |
| *Telmatoscopus - Trichomyia* | 1.1057 | 0.526 | 115 | 2.102 | 0.5783 |

**TABLE S5.** Tukey tests for PC2.

| **contrast** | **estimate** | **SE** | **df** | **t.ratio** | **p.value** |
|---|---|---|---|---|---|
| *Brumptomyia - Clytocerus* | 0.039646 | 0.647 | 115 | 0.061 | 1 |
| *Brumptomyia - Lutzomyia* | 0.228164 | 0.506 | 115 | 0.451 | 1 |
| *Brumptomyia - Pericoma* | 0.323603 | 0.571 | 115 | 0.567 | 1 |
| *Brumptomyia - Philosepedon* | -0.000583 | 0.737 | 115 | -0.001 | 1 |
| *Brumptomyia - Phlebotomus* | -0.592711 | 0.571 | 115 | -1.039 | 0.9938 |
| *Brumptomyia - Psychoda* | 0.269408 | 0.522 | 115 | 0.516 | 1 |
| *Brumptomyia - Satchelliella* | 0.288124 | 0.571 | 115 | 0.505 | 1 |
| *Brumptomyia - Sergentomyia* | 0.203861 | 0.58 | 115 | 0.352 | 1 |
| *Brumptomyia - Telmatoscopus* | -0.022894 | 0.58 | 115 | -0.04 | 1 |
| *Brumptomyia - Trichomyia* | -0.380886 | 0.737 | 115 | -0.517 | 1 |
| *Clytocerus - Lutzomyia* | 0.188519 | 0.457 | 115 | 0.412 | 1 |
| *Clytocerus - Pericoma* | 0.283957 | 0.528 | 115 | 0.538 | 1 |
| *Clytocerus - Philosepedon* | -0.040229 | 0.704 | 115 | -0.057 | 1 |

| | | | | | |
|---|---|---|---|---|---|
| *Clytocerus - Phlebotomus* | -0.632357 | 0.528 | 115 | -1.197 | 0.9816 |
| *Clytocerus - Psychoda* | 0.229762 | 0.476 | 115 | 0.483 | 1 |
| *Clytocerus - Satchelliella* | 0.248478 | 0.528 | 115 | 0.47 | 1 |
| *Clytocerus - Sergentomyia* | 0.164215 | 0.538 | 115 | 0.305 | 1 |
| *Clytocerus - Telmatoscopus* | -0.06254 | 0.538 | 115 | -0.116 | 1 |
| *Clytocerus - Trichomyia* | -0.420532 | 0.704 | 115 | -0.597 | 0.9999 |
| *Lutzomyia - Pericoma* | 0.095438 | 0.341 | 115 | 0.28 | 1 |
| *Lutzomyia - Philosepedon* | -0.228748 | 0.577 | 115 | -0.396 | 1 |
| *Lutzomyia - Phlebotomus* | -0.820875 | 0.341 | 115 | -2.408 | 0.3712 |
| *Lutzomyia - Psychoda* | 0.041243 | 0.252 | 115 | 0.163 | 1 |
| *Lutzomyia - Satchelliella* | 0.059959 | 0.341 | 115 | 0.176 | 1 |
| *Lutzomyia - Sergentomyia* | -0.024304 | 0.356 | 115 | -0.068 | 1 |
| *Lutzomyia - Telmatoscopus* | -0.251058 | 0.356 | 115 | -0.706 | 0.9998 |
| *Lutzomyia - Trichomyia* | -0.60905 | 0.577 | 115 | -1.055 | 0.993 |
| *Pericoma - Philosepedon* | -0.324186 | 0.635 | 115 | -0.511 | 1 |
| *Pericoma - Phlebotomus* | -0.916314 | 0.431 | 115 | -2.125 | 0.5625 |
| *Pericoma - Psychoda* | -0.054195 | 0.365 | 115 | -0.148 | 1 |
| *Pericoma - Satchelliella* | -0.035479 | 0.431 | 115 | -0.082 | 1 |
| *Pericoma - Sergentomyia* | -0.119742 | 0.443 | 115 | -0.27 | 1 |
| *Pericoma - Telmatoscopus* | -0.346497 | 0.443 | 115 | -0.782 | 0.9994 |
| *Pericoma - Trichomyia* | -0.704489 | 0.635 | 115 | -1.11 | 0.9896 |
| *Philosepedon - Phlebotomus* | -0.592128 | 0.635 | 115 | -0.933 | 0.9974 |
| *Philosepedon - Psychoda* | 0.269991 | 0.592 | 115 | 0.456 | 1 |
| *Philosepedon - Satchelliella* | 0.288707 | 0.635 | 115 | 0.455 | 1 |

| | | | | | |
|---|---|---|---|---|---|
| *Philosepedon - Sergentomyia* | 0.204444 | 0.643 | 115 | 0.318 | 1 |
| *Philosepedon - Telmatoscopus* | -0.022311 | 0.643 | 115 | -0.035 | 1 |
| *Philosepedon - Trichomyia* | -0.380303 | 0.787 | 115 | -0.483 | 1 |
| *Phlebotomus - Psychoda* | 0.862118 | 0.365 | 115 | 2.36 | 0.4015 |
| *Phlebotomus - Satchelliella* | 0.880835 | 0.431 | 115 | 2.042 | 0.6201 |
| *Phlebotomus - Sergentomyia* | 0.796571 | 0.443 | 115 | 1.798 | 0.7791 |
| *Phlebotomus - Telmatoscopus* | 0.569817 | 0.443 | 115 | 1.286 | 0.9695 |
| *Phlebotomus - Trichomyia* | 0.211825 | 0.635 | 115 | 0.334 | 1 |
| *Psychoda - Satchelliella* | 0.018716 | 0.365 | 115 | 0.051 | 1 |
| *Psychoda - Sergentomyia* | -0.065547 | 0.379 | 115 | -0.173 | 1 |
| *Psychoda - Telmatoscopus* | -0.292301 | 0.379 | 115 | -0.771 | 0.9995 |
| *Psychoda - Trichomyia* | -0.650293 | 0.592 | 115 | -1.098 | 0.9904 |
| *Satchelliella - Sergentomyia* | -0.084263 | 0.443 | 115 | -0.19 | 1 |
| *Satchelliella - Telmatoscopu* | s -0.311018 | 0.443 | 115 | -0.702 | 0.9998 |
| *Satchelliella - Trichomyia* | -0.66901 | 0.635 | 115 | -1.054 | 0.993 |
| *Sergentomyia - Telmatoscopus* | -0.226754 | 0.455 | 115 | -0.499 | 1 |
| *Sergentomyia - Trichomyia* | -0.584747 | 0.643 | 115 | -0.91 | 0.9979 |
| *Telmatoscopus - Trichomyia* | -0.357992 | 0.643 | 115 | -0.557 | 1 |

**TABLE S6.** Tukey tests for PC3.

| contrast | estimate | SE | df | t.ratio | p.value |
|---|---|---|---|---|---|
| *Brumptomyia - Clytocerus* | 0.0496 | 0.51 | 115 | 0.097 | 1 |
| *Brumptomyia - Lutzomyia* | -0.3483 | 0.398 | 115 | -0.874 | 0.9985 |

| | | | | | |
|---|---|---|---|---|---|
| *Brumptomyia - Pericoma* | -0.0169 | 0.449 | 115 | -0.038 | 1 |
| *Brumptomyia - Philosepedon* | -0.0968 | 0.58 | 115 | -0.167 | 1 |
| *Brumptomyia - Phlebotomus* | 1.1306 | 0.449 | 115 | 2.515 | 0.3069 |
| *Brumptomyia - Psychoda* | 0.0853 | 0.412 | 115 | 0.207 | 1 |
| *Brumptomyia - Satchelliella* | 0.1567 | 0.449 | 115 | 0.349 | 1 |
| *Brumptomyia - Sergentomyia* | 1.6562 | 0.457 | 115 | 3.628 | 0.0178 |
| *Brumptomyia - Telmatoscopus* | -0.2788 | 0.457 | 115 | -0.611 | 0.9999 |
| *Brumptomyia - Trichomyia* | 0.2274 | 0.58 | 115 | 0.392 | 1 |
| *Clytocerus - Lutzomyia* | -0.3979 | 0.36 | 115 | -1.104 | 0.99 |
| *Clytocerus - Pericoma* | -0.0665 | 0.416 | 115 | -0.16 | 1 |
| *Clytocerus - Philosepedon* | -0.1464 | 0.555 | 115 | -0.264 | 1 |
| *Clytocerus - Phlebotomus* | 1.081 | 0.416 | 115 | 2.598 | 0.2624 |
| *Clytocerus - Psychoda* | 0.0357 | 0.375 | 115 | 0.095 | 1 |
| *Clytocerus - Satchelliella* | 0.1071 | 0.416 | 115 | 0.257 | 1 |
| *Clytocerus - Sergentomyia* | 1.6066 | 0.424 | 115 | 3.791 | 0.0105 |
| *Clytocerus - Telmatoscopus* | -0.3284 | 0.424 | 115 | -0.775 | 0.9995 |
| *Clytocerus - Trichomyia* | 0.1778 | 0.555 | 115 | 0.321 | 1 |
| *Lutzomyia - Pericoma* | 0.3314 | 0.269 | 115 | 1.234 | 0.9772 |
| *Lutzomyia - Philosepedon* | 0.2515 | 0.455 | 115 | 0.553 | 1 |
| *Lutzomyia - Phlebotomus* | 1.4789 | 0.269 | 115 | 5.505 | <.0001 |
| *Lutzomyia - Psychoda* | 0.4336 | 0.199 | 115 | 2.181 | 0.5231 |
| *Lutzomyia - Satchelliella* | 0.505 | 0.269 | 115 | 1.88 | 0.7288 |
| *Lutzomyia - Sergentomyia* | 2.0045 | 0.28 | 115 | 7.151 | <.0001 |
| *Lutzomyia - Telmatoscopus* | 0.0695 | 0.28 | 115 | 0.248 | 1 |

| | | | | | |
|---|---|---|---|---|---|
| *Lutzomyia - Trichomyia* | 0.5757 | 0.455 | 115 | 1.266 | 0.9727 |
| *Pericoma - Philosepedon* | -0.0799 | 0.5 | 115 | -0.16 | 1 |
| *Pericoma - Phlebotomus* | 1.1475 | 0.34 | 115 | 3.377 | 0.0383 |
| *Pericoma - Psychoda* | 0.1022 | 0.288 | 115 | 0.355 | 1 |
| *Pericoma - Satchelliella* | 0.1737 | 0.34 | 115 | 0.511 | 1 |
| *Pericoma - Sergentomyia* | 1.6731 | 0.349 | 115 | 4.793 | 0.0003 |
| *Pericoma - Telmatoscopus* | -0.2619 | 0.349 | 115 | -0.75 | 0.9996 |
| *Pericoma - Trichomyia* | 0.2444 | 0.5 | 115 | 0.489 | 1 |
| *Philosepedon - Phlebotomus* | 1.2274 | 0.5 | 115 | 2.454 | 0.3427 |
| *Philosepedon - Psychoda* | 0.1821 | 0.466 | 115 | 0.39 | 1 |
| *Philosepedon - Satchelliella* | 0.2535 | 0.5 | 115 | 0.507 | 1 |
| *Philosepedon - Sergentomyia* | 1.753 | 0.507 | 115 | 3.461 | 0.0299 |
| *Philosepedon - Telmatoscopus* | -0.182 | 0.507 | 115 | -0.359 | 1 |
| *Philosepedon - Trichomyia* | 0.3242 | 0.62 | 115 | 0.523 | 1 |
| *Phlebotomus - Psychoda* | -1.0453 | 0.288 | 115 | -3.632 | 0.0176 |
| *Phlebotomus - Satchelliella* | -0.9738 | 0.34 | 115 | -2.866 | 0.1474 |
| *Phlebotomus - Sergentomyia* | 0.5256 | 0.349 | 115 | 1.506 | 0.9151 |
| *Phlebotomus - Telmatoscopus* | -1.4094 | 0.349 | 115 | -4.037 | 0.0045 |
| *Phlebotomus - Trichomyia* | -0.9032 | 0.5 | 115 | -1.806 | 0.7743 |
| *Psychoda - Satchelliella* | 0.0714 | 0.288 | 115 | 0.248 | 1 |
| *Psychoda - Sergentomyia* | 1.5709 | 0.299 | 115 | 5.259 | <.0001 |
| *Psychoda - Telmatoscopus* | -0.3641 | 0.299 | 115 | -1.219 | 0.9791 |
| *Psychoda - Trichomyia* | 0.1421 | 0.466 | 115 | 0.305 | 1 |
| *Satchelliella - Sergentomyia* | 1.4994 | 0.349 | 115 | 4.295 | 0.0018 |

| | | | | | |
|---|---|---|---|---|---|
| *Satchelliella - Telmatoscopus* | -0.4355 | 0.349 | 115 | -1.248 | 0.9753 |
| *Satchelliella - Trichomyia* | 0.0707 | 0.5 | 115 | 0.141 | 1 |
| *Sergentomyia - Telmatoscopus* | -1.935 | 0.358 | 115 | -5.403 | <.0001 |
| *Sergentomyia - Trichomyia* | -1.4287 | 0.507 | 115 | -2.821 | 0.1636 |
| *Telmatoscopus - Trichomyia* | 0.5062 | 0.507 | 115 | 0.999 | 0.9954 |

**TABLE S7. Trait-evolution models suggest that PC1—a proxy of climatic niche evolution— in Psychodidae evolves according to a Brownian motion model of trait evolution.**

| model | Parameters | AIC | wAIC |
|---|---|---|---|
| Early burst | a = -1.28, sigsq = 3.98, z0 = -0.89 | 155.39 | 0.15 |
| Brownian motion | sigsq = 2.66, z0 = -0.91 | 153.62 | 0.35 |
| Ornstein–Uhlenbeck | alpha = 0.00, sigsq = 2.66, z0 = -0.91 | 155.79 | 0.12 |
| Delta | delta = 0.76, sigsq = 3.05, z0 = -0.90 | 155.55 | 0.14 |
| Lambda | lambda = 1.00, sigsq = 2.66, z0 = -0.91 | 155.79 | 0.12 |
| White-noise | sigsq = 2.19, z0 = -0.59 | 272.08 | 0.00 |
| kappa | kappa = 0.89, sigsq = 2.00, z0 = -0.87 | 155.68 | 0.13 |

**TABLE S8. Trait-evolution models suggest that mean latitude proxy of climatic niche evolution— in Psychodidae evolves according to a Early Burst model of trait evolution.**

| model | Parameters | AIC | aic.weights |
|---|---|---|---|
| Early burst | a = -6.54, sigsq = 9175.83, z0 = 0.02 | 606.42 | 0.82 |
| Brownian motion | sigsq = 1313.68, z0 = -0.38 | 612.72 | 0.04 |
| Ornstein–Uhlenbeck | alpha = 0.00, sigsq = 1313.19, z0 = -0.38 | 614.89 | 0.01 |
| Delta | delta = 0.26, sigsq = 3134.84, z0 = -0.82 | 610.31 | 0.12 |
| Lambda | lambda = 1.00, sigsq = 1313.68, z0 = -0.38 | 614.89 | 0.01 |
| White-noise | sigsq = 745.36, z0 = 17.92 | 703.60 | 0.00 |
| kappa | kappa = 1.00, sigsq = 1313.68, z0 = -0.38 | 614.89 | 0.01 |

**FIGURE S1. Geographic distribution of the samples included in this study. A.** Location of the included samples. **B.** Mean latitude distribution for all species in the Psychodidae family. The solid black line marks the equator; dashed black lines mark the boundaries of the tropics.

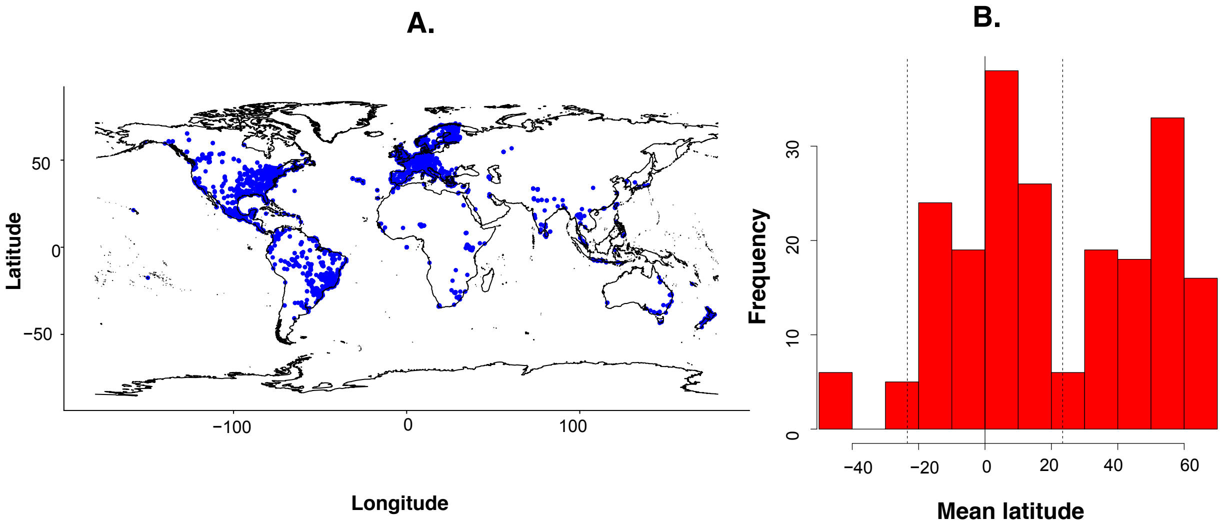

**FIGURE S2.** Mean latitude distribution for 72 species from 12 genera in the Psychodidae family. All species had at least five georeferenced records. Solid black line marks the equator; dashed black lines mark the boundaries of the tropics. Purple bars represent non-hematophagous species, orange bars represent hematophagous species who do not carry *Leishmania*. Purple bars represent hematophagous vector species.

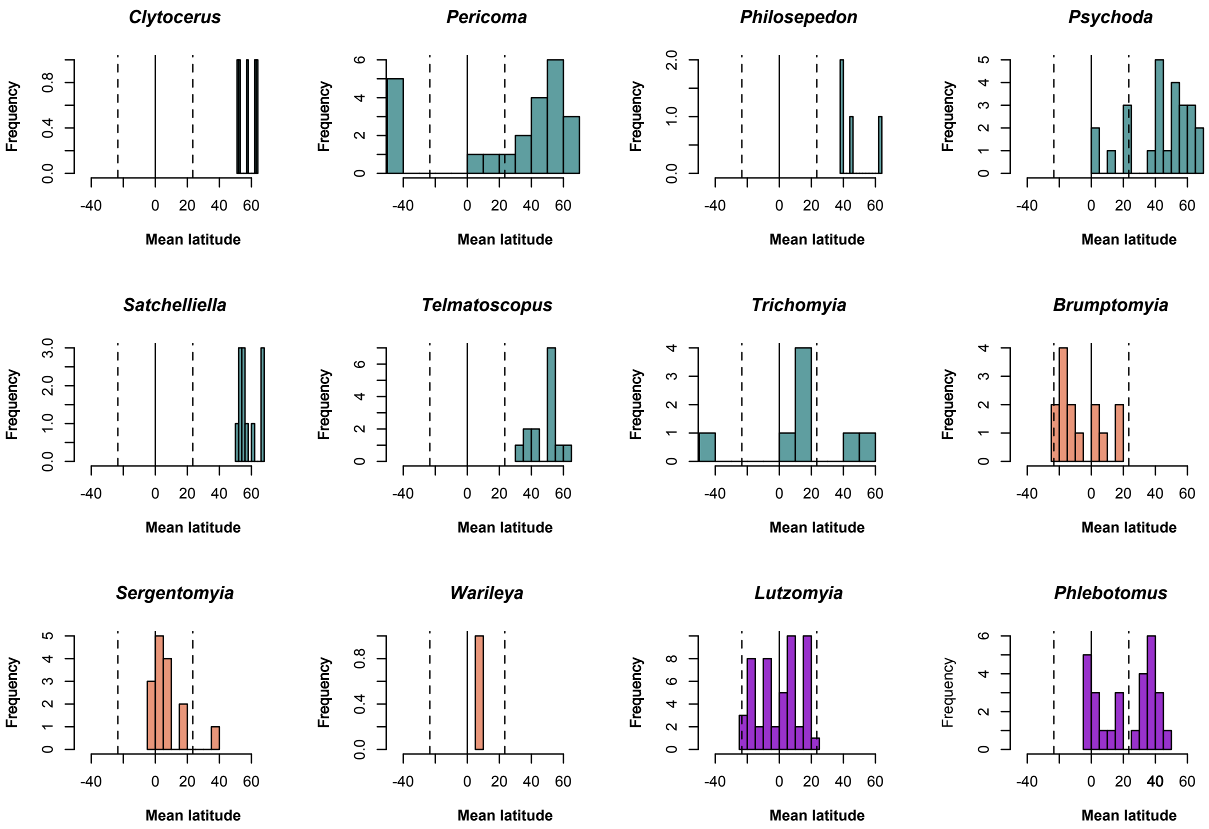

**FIGURE S3.** *COI* gene genealogy for the Psychodidae family. Values on each node show the bootstrap support with 1,000 replicates.

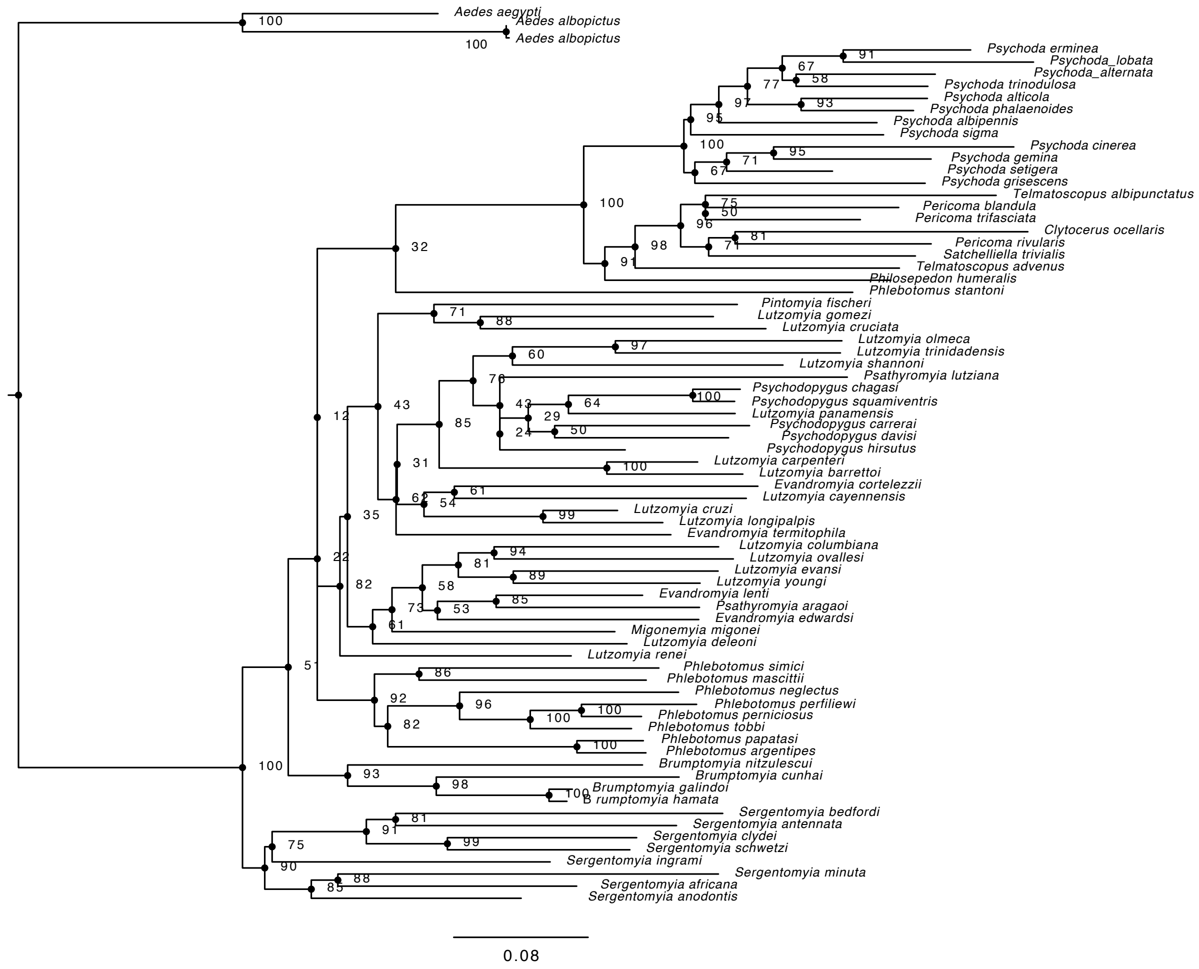

**FIGURE S4. Ancestral trait reconstruction using three different proxies of geographical distribution values suggest the ancestor of the Psychodidae family had a climatic niche akin to a tropical species. A.** PC1 of climatic variation. **B.** Mean latitude. Figure 4 shows the ancestral niche reconstructions for Tropicality index.

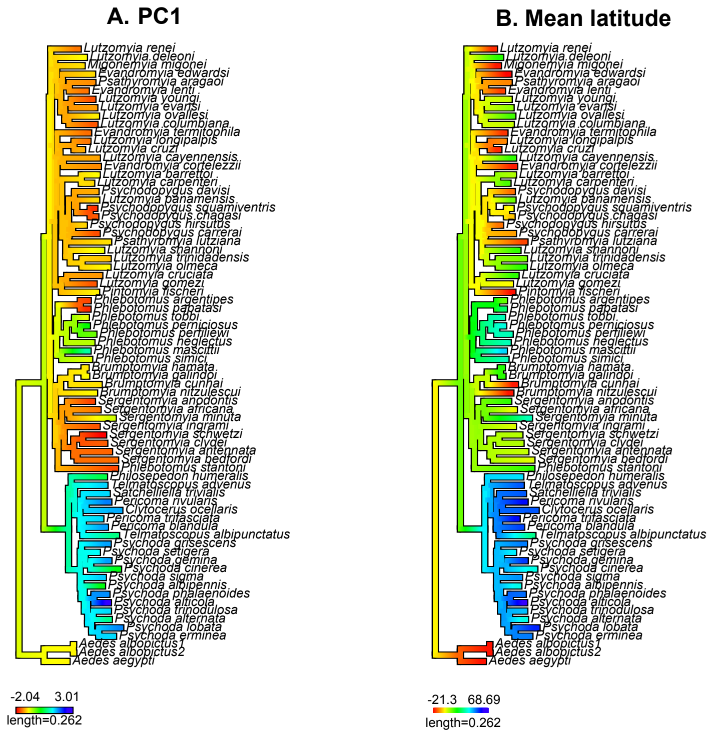
